# Efficient *in vitro* refactoring and biosynthetic gene cluster amplification for the overproduction and accelerated discovery of anticancer thioamitides

**DOI:** 10.64898/2026.01.07.698260

**Authors:** Javier Santos-Aberturas, Luca Frattaruolo, Eleonora Tassinari, Martin Rejzek, Louis K. W. Fong, Praphat Kawicha, Aphidech Sangdee, Carolina Grandellis, Anna Rita Cappello, Andrew W. Truman

## Abstract

Thioamitides, a class of highly modified bacterial ribosomally synthesised and post-translationally modified peptides (RiPPs), have potent activities against multiple cancer cell lines. Among these compounds, the structurally divergent thioalbamide combines promising *in vivo* antiproliferative activity with a superior chemical stability respect to its counterparts. However, thioalbamide is produced in low yields by its genetically intractable native producer and its biosynthetic pathway was initially not productive when transferred into the heterologous host *Streptomyces coelicolor* M1146. These circumstances substantially hamper to increase the production of this promising compound. Here, we show how *in vitro* Gibson-like assemblies can be employed for the quick and efficient refactoring of the thioalbamide biosynthetic gene cluster (BGC), leading to substantially increased levels of production in *S. coelicolor* M1146 through a prioritised selection of promoters. Via this work, P*tsrA* and P*groEL2* were identified as beneficial additions to the *Streptomyces* synthetic biology toolbox. We then assessed bacterial genomes for biosynthetic gene clusters (BGCs) predicted to produce thioalbamide-like compounds with improved hydrophilicity. This rational discovery campaign led to the identification a silent thioamitide BGC encoding a thioalbamide-like core peptide but clustered with additional tailoring enzymes, including a previously unknown cupin-fold protein. Applying the refactoring strategy together with the simultaneous expression of multiple BGC copies, we characterised the product of this pathway, thiocupinamide, a polyhydroxylated thioamitide closely related to thioalbamide. We show that thiocupinamide has potent anticancer and antibacterial activities.

## INTRODUCTION

Thioamitides constitute a class of bacterial ribosomally synthesised and post-translationally modified peptides (RiPPs) that are biosynthetically related to the recently established class V lanthipeptides^1–3^. These compounds are among the most complex RiPPs, with their general structure (**Figure 1**) including a lactyl or pyruvyl group at their N-terminus, a linear region including between three and five thioamide bonds, and a C-terminal macrocycle closed by an aminovinylcysteine moiety^4–6^. Within this macrocycle there is an N,N-dimethylated histidinium residue that confers a permanent positive charge and is conserved across all characterised thioamitides. Thioamitides have promising anticancer properties as they induce apoptosis through the inhibition of the mitochondrial F_1_-F_0_ ATP synthase complex, and have therefore attracted substantial research interest since the discovery of their first representative, thioviridamide^7–11^.

**Figure 1.**
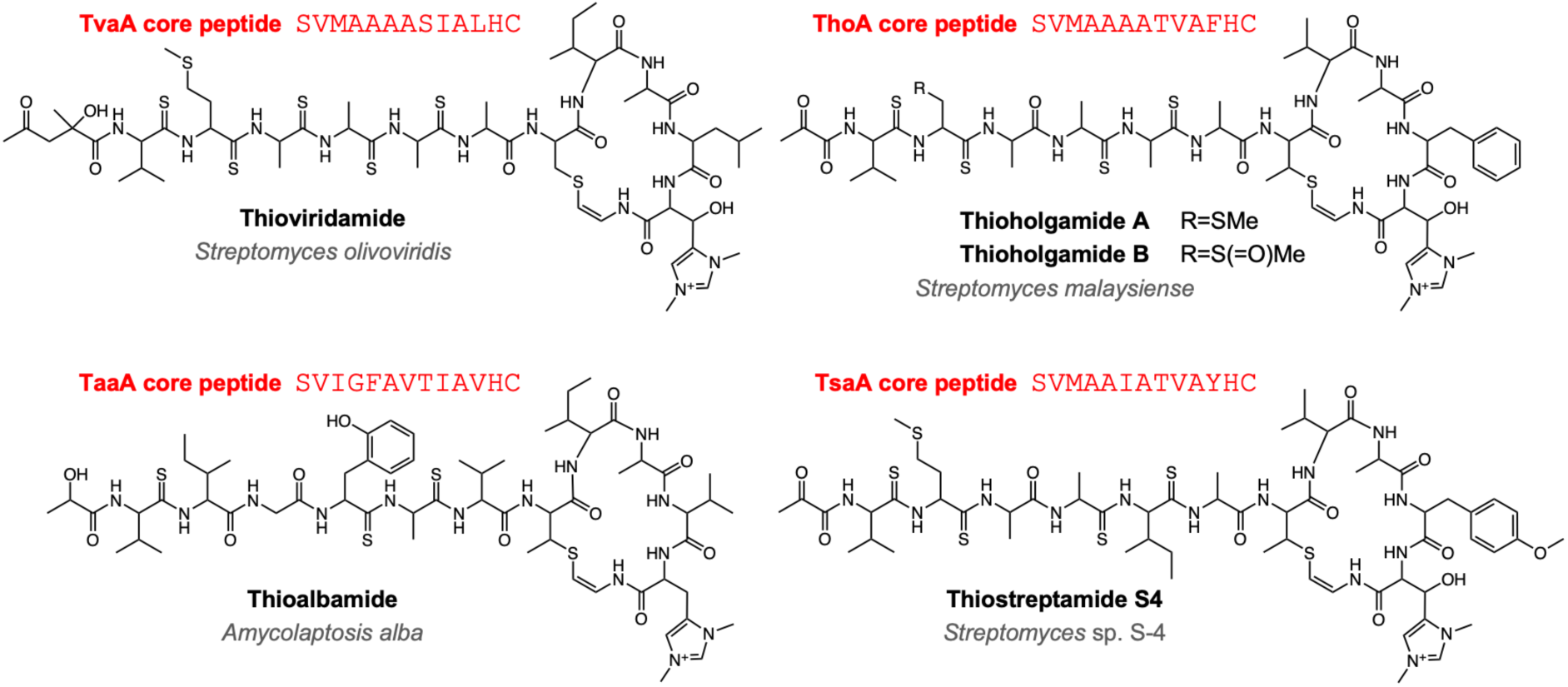
Examples of thioamitide structures and associated core peptides.

Following the identification of the thioviridamide biosynthetic gene cluster (BGC)^12,13^, genome mining approaches have led to the discovery of several new thioamitides, partially unveiling the structural diversity of these RiPPs^4,6,14^. The general biosynthetic pathway of these compounds is well understood, thanks to the combination of genetic, metabolomic and biochemical methods^15–20^ (**Figure 2**). Prior studies have shown that thioamitide biosynthetic machinery is partially tolerant to the introduction of amino acid changes within the core peptide, thus facilitating the creation of libraries to be screened in the search for variants displaying improved pharmacological properties^20–22^.

**Figure 2.**
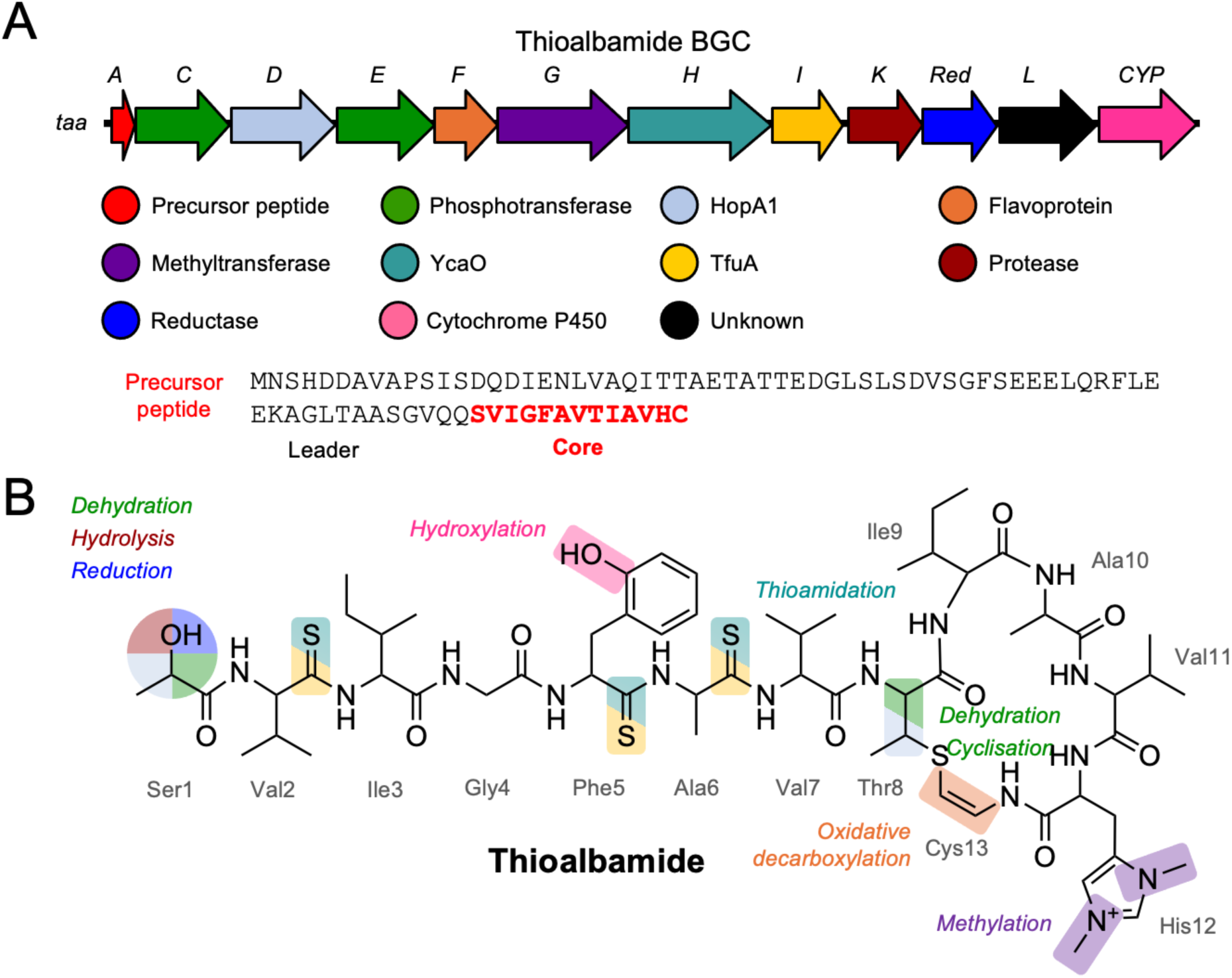
Thioalbamide BGC and chemical structure. (A) Thioalbamide BGC. Protein functions are colour-coded. The sequence of the precursor peptide is shown, with the core peptide highlighted in red. (B) Thioalbamide structure. The chemical moieties introduced as the result of post-translational modifications of the core peptide are highlighted with colours in accordance with the genes indicated in panel A.

Thioalbamide (**Figure 1**) is the most structurally divergent thioamitide, and an interesting drug lead candidate due to its activity in suppressing tumour proliferation in a mouse model^8^, as well as having some of the most potent *in vitro* cytotoxicity towards cancer cells among currently characterised thioamitides^6,23^. Unlike most of its counterparts (**Figure 1**), in which the linear parts of these molecule are rich in alanine residues, with four or five thioamide bonds, thioalbamide contains a variety of residues and only three thioamidations. Additionally, thioalbamide lacks a β-hydroxylation of the N,N-dimethylated histidinium, which is present in all other thioamitides. However, the potentially most important difference is the third residue of the core peptide. This residue is a methionine in all characterised natural thioamitides except in thioalbamide, where it is substituted by an isoleucine. Although all thioamitides exhibit potent cytotoxic activities in the nanomolar range, the spontaneous oxidation of the sulfur atom of Met3 leads to a 10-fold diminishment in their biological activity^4^. Thus, thioalbamide is more stable to oxidation than most other thioamitides, and therefore more likely to constitute a suitable drug lead.

However, several constraints have limited a deeper exploration of the biological properties of thioalbamide and its therapeutic potential. First, only low yields of this compound can be obtained from complex solid medium cultures of its native producer, *Amycolatopsis alba* DSM 44262. Second, we have not been able to genetically manipulate *A. alba*, which precludes efforts to improve production via strain engineering. Third, and unlike in the case of other thioamitides^13,14,20,22^, the introduction of the native thioalbamide BGC in standard *Streptomyces* heterologous hosts does not result in the production of the compound. Fourth, and despite its promising bioactivity and chemical stability, the high hydrophobicity of thioalbamide may not be ideal for drug development and formulation.

Here, to overcome these challenges we report the development of a quick *in vitro* refactoring strategy for the heterologous overproduction of thioalbamide in *Streptomyces coelicolor* M1146^24^, including the identification of P*tsrA* as a new promising promoter for *Streptomyces* synthetic biology, and we show how the integration of additional pathway copies can increase production levels. We apply these principles to accelerate the targeted discovery and characterization of thiocupinamide, a new thioalbamide analogue featuring three additional hydroxylations, two of them introduced on the key Ile3 residue through the action of a novel cupin tailoring enzyme. Thiocupinamide has potent cytotoxic activity towards numerous cancer cell lines and improved antimicrobial activity compared to thioalbamide.

## RESULTS AND DISCUSSION

### Heterologous overproduction of thioalbamide in *S. coelicolor* M1146

We initially employed transformation-associated recombination (TAR) cloning in yeast to capture the thioalbamide BGC of *A. alba* DSM 44262 into pCAP03. However, and unlike in the case of the thiostreptamide S4 BGC, the introduction of this construct (pCAP03_*taa*_BGC) into *S. coelicolor* M1146 did not result in the production of thioalbamide. A gene encoding an AfsR/SARP regulator^25^ adjacent to *taaA* was a potential candidate as a pathway-specific regulator, but was not originally captured in pCAP03_alba_BGC. Thus, we generated a compatible construct (pLF026) for the constitutive expression of this regulator and conjugated it into *S. coelicolor* M1146 pCAP03_*taa*_BGC. Disappointingly, this combination did not lead to the production of thioalbamide, suggesting that the AfsR/SARP regulator is not sufficient by itself to trigger the transcription of the thioalbamide BGC, or simply that this gene is not associated with thioalbamide regulation.

Given these initial difficulties, we opted for a refactoring strategy, taking advantage of the unusually simple transcriptional structure of the thioalbamide BGC (**Figure 2A**), in which all the biosynthetic genes are oriented in the same direction and are overlapping or with very short intergenic spaces. This organization includes the precursor peptide coding gene *taa*A, which is separated by only 12 bp. This strongly suggests the lack of additional regulatory elements in this intergenic region, and that the whole BGC might be transcribed as a single transcript, notably simplifying the refactoring strategy to trigger its expression. It is important to notice that given the structural role of the precursor peptide in RiPP biosynthetic pathways, its expression levels constitute the first potential bottleneck for RiPP production levels and strong transcription and translation are intuitively beneficial for their yields. Therefore, we employed Gibson-like isothermal *in vitro* assemblies to generate a library of refactored thioalbamide BGCs under the control of a curated selection of ten promoters (**Table S3**, **Figure 3**), including established *Streptomyces* constitutive promoters and RiPP promoters known to work in *S. coelicolor* M1146 (P*ermE\**, P*btmC*, P*kasO\**, PSF14, P*tsaA*, P*accIV*, P*varA* and P*hrdB*)^26–28^, alongside some untested promoters that we considered potentially interesting based on existing literature, such as P*groEL2* and P*tsrA*^29,30^. *In vitro* assembly of PCR products proved to be more rapid that TAR cloning for a BGC of this size, which constituted a four-part assembled of two BGC fragments, the variable promoter fragment and the pSET152 backbone^31^, which features the φC31 integrase. These constructs were introduced in *S. coelicolor* M1146 through a semi-automated robotic workflow for conjugation. Here, *Escherichia coli* transformation and *E. coli–Streptomyces* conjugation steps were carried out using a series of automated robotic systems (see Materials and Methods and **Figure S1**). Exconjugants were subsequently manually selected, cultured, and progressed to downstream analysis. This workflow demonstrates the potential for future scalability to support larger screening efforts. The integration of robotic liquid handlers increases reproducibility, reduces experimental error, and minimises time-consuming tasks such as pipetting and plating.

**Figure 3.**
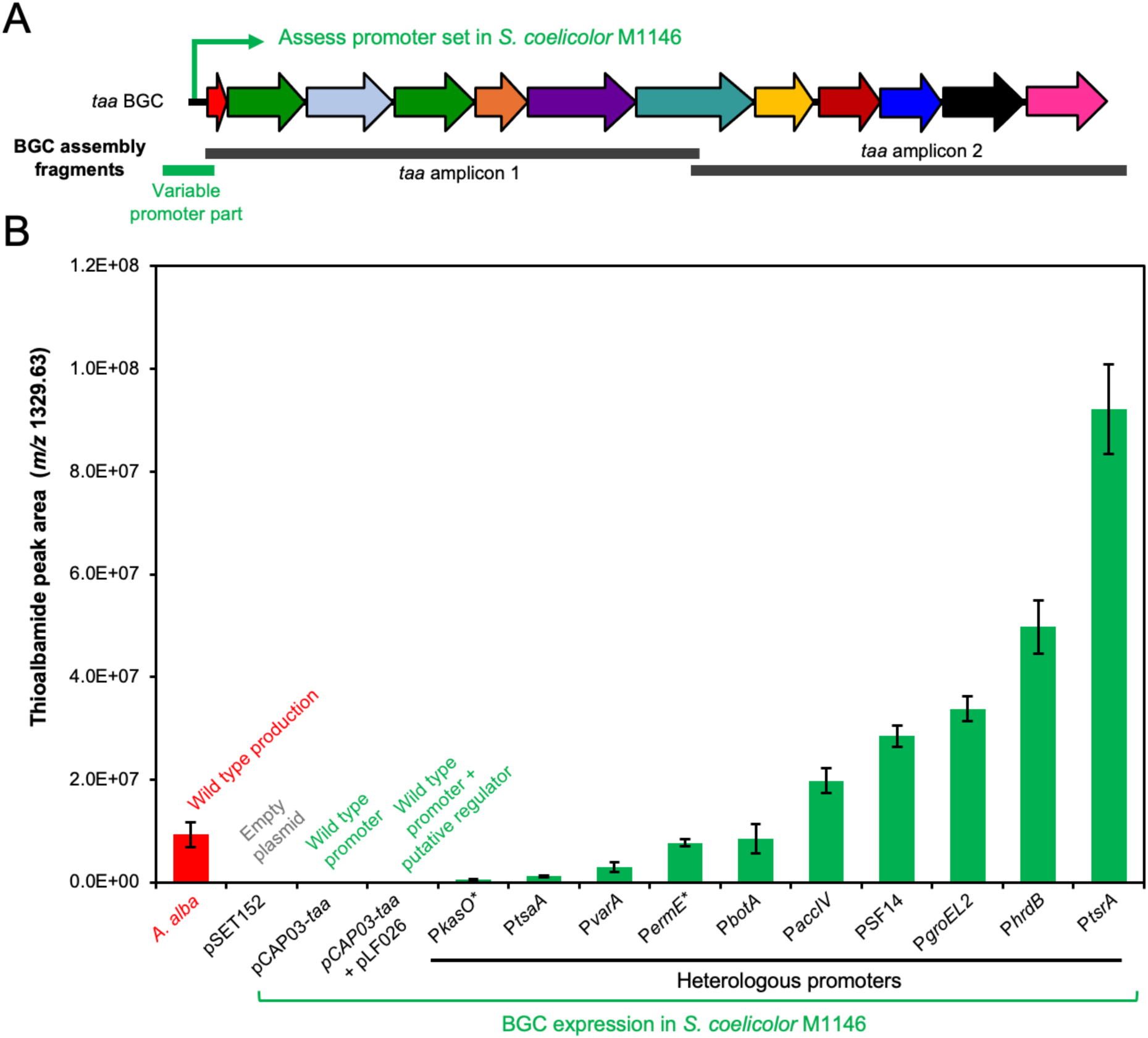
Thioalbamide BGC refactoring triggers heterologous production in *S. coelicolor* M1146. (A) Overview of refactoring strategy for cloning into pSET152. (B) Thioalbamide production assessed by LC-MS. The mean values of extracted ion chromatogram peak areas (*m/z* 1329.63) from LC-MS analysis of four fermentation replicates are shown, with vertical bars representing standard deviation. As a comparison, wild type production provides approximately 2 mg/L purified product.

Once integrated into the genome of *S. coelicolor* M1146, all these constructs led to the production of thioalbamide, although with very diverse yields (**Figure 3B**) when assessed using liquid chromatography - mass spectrometry (LC-MS). Interestingly, some of the established *Streptomyces* promoters perceived as strong and constitutive (such as P*ermE\** or P*kasO\**) exhibited relatively poor performances, while the highest yields were obtained with P*tsrA*, which achieved a seven-fold production increase respect to the yields detected in *A. alba*. P*tsrA* is the promoter that drives the expression of the precursor peptide of the thiostrepton pathway in *Streptomyces laurentii*, which one of the most productive RiPP pathways reported, where thiostrepton yields of >10 g/L were reported from fed-batch cultures^32^. In light of these results, P*tsrA* could represent a useful addition to the *Streptomyces* synthetic biology toolbox.

### Discovery and heterologous expression of the thiocupinamide BGC

Once we established a strategy for the heterologous overproduction of thioalbamide, we decided to explore its application to the discovery of new thioalbamide analogues with increased hydrophilicity. We therefore conducted targeted genome mining using a RiPPER-based workflow^28,33^ to obtain an updated view of thioamitide BGC diversity. Here, candidate BGCs were identified using HopA1 proteins as bait for RiPPER, which was used to retrieve the corresponding precursor peptides (see Materials and Methods for full workflow details). Sequence identity networking was used to compare precursor peptide sequences (**Figure 4A** and **Figure S2**), which revealed distinct core peptide sequences motifs that we propose can be sub-divided into four subfamilies. The associated BGCs each encode proteins expected for thioamitide biosynthesis, as well as further enzymes that are predicted to introduce further diversity into this RiPP family (**Figure 4B, Figures S3-S5**).

**Figure 4.**
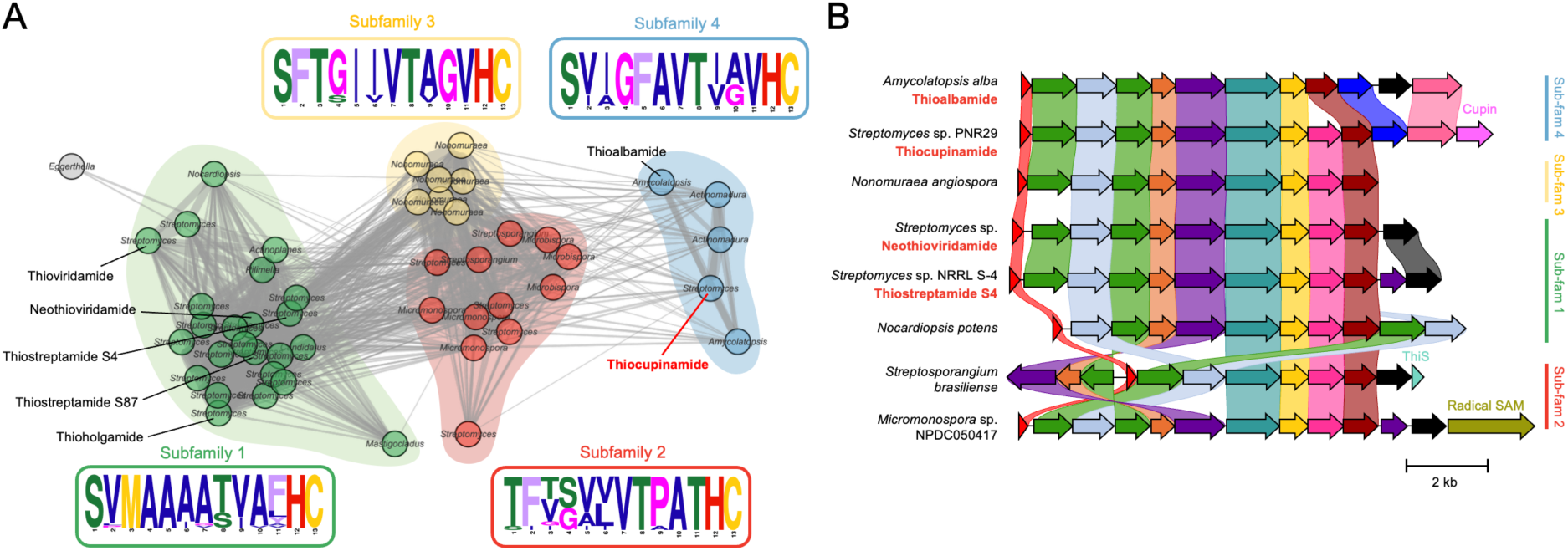
Mapping the diversity of thioamitide precursor peptides and BGCs. (A) Precursor peptide sequence identity networking reveals subfamilies of thioamitides based on core similarity. Network edges are shown for peptides with over 40% sequence identity and at least 40% sequence coverage. Nodes are colour-coded based on subfamily, while sequence logos are colour-coded based on amino acid chemistry (see Materials and Methods for full details). Characterised precursor peptides are annotated. (B) Example BGCs corresponding to each subfamily. Genes links are shown if sequence identity >30%.

Notably, almost all characterised thioamitides come from subfamily 1, which is the only subfamily to feature the oxidation-sensitive Met3. TaaA belongs to subfamily 4, which features related precursor peptides with either isoleucine or alanine at position 3 (**Figure 4A**). The corresponding BGCs were assessed for tailoring enzymes with functionalities that could introduce polar moieties in the final products. This analysis revealed that, unlike the thioalbamide BGC, several of these BGCs include a gene encoding a dioxygenase (pfam05721) homologous to hydroxylases that are responsible for the ý-hydroxylation of the N,N-dimethyl histidinium moiety in other thioamitides^17^ (**Figure 1**). In addition, some BGCs included an intriguing additional gene encoding a cupin superfamily protein (pfam08007) with no homologues in any previously characterized thioamitide BGCs. Cupins are an extremely versatile superfamily of proteins^34^, and the enzymes among them can catalyse both halogenations and oxygenations that sometimes lead to complex rearrangements in different specialized metabolism pathways^35^.

To unveil the final product of one representative of this new subset of thioamitide biosynthetic pathways, we focused our efforts on a candidate BGC **(Figure 5A, Table S4)** within the genome of *Streptomyces* sp. S.PNR29, a strain with biocontrol properties isolated in Thailand^36^. Due to the presence of the novel cupin enzyme encoded in the BGC, we putatively named it the thiocupinamide (*tca*) BGC. The only difference in the core peptide compared to thioalbamide is a Val9 residue instead of Ile9, suggesting a less hydrophobic peptide backbone. The *tca* BGC displays a similar transcriptional organization to the *taa* BGC, with an intergenic space of just 13 bp between *tcaA* and *tcaC*. Following the strategy described above, we refactored the *tca* BGC as a single transcriptional unit under the control of P*tsrA*, using pCAP03^37^ as vector backbone (pCAP03_P*tsrA*_*tca*_BGC). To understand the role of the novel cupin (TcaO), we also obtained a refactored version of the cluster lacking its coding gene (pCAP03_P*tsrA*_ *tca*_BGC_Δ*tcaO*).

**Figure 5.**
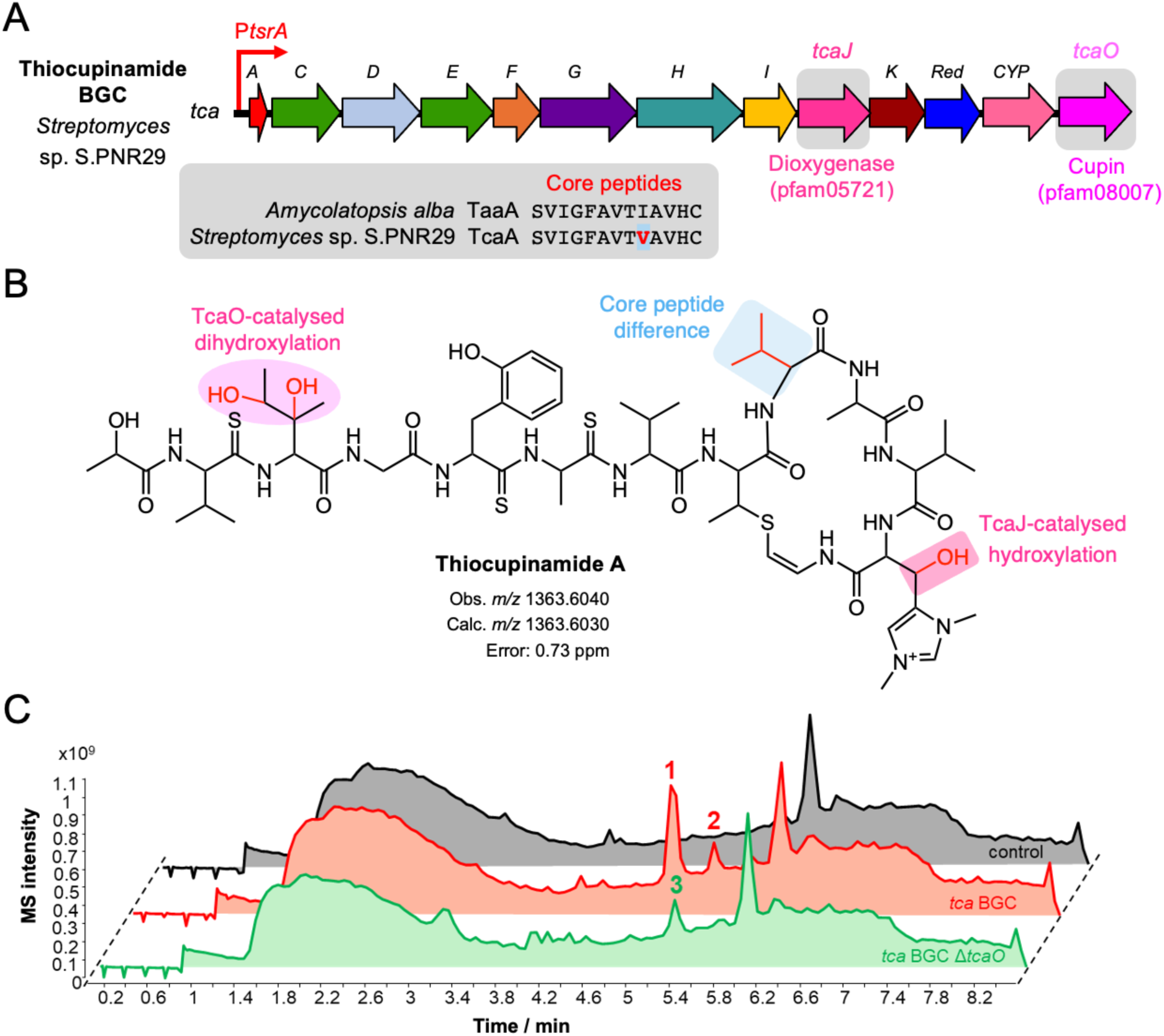
Thiocupinamide discovery. (A) Thiocupinamide (*tca*) BGC. Gene functions are colour-coded as in Figure 2A. Genes are named based on the thioalbamide BGC. Additional genes with respect to the thioalbamide BGC are highlighted, along with the difference in core peptide sequence. (B) Thiocupinamide A structure, highlighting the structural differences compared to thioalbamide. (C) Total ion chromatograms showing thiocupinamide production by heterologous expression in *S. coelicolor* M1146. TcaO introduces two oxygen atoms into thiocupinamide precursors to generate the final product of the pathway, thiocupinamide A (**1**). Note the differential peaks produced by *S. coelicolor* M1146 with pSET152 P*tsrA tca* BGC (red) or pSET152 P*tsrA tca* BGC Δ*tcaO* (green) compared to a control strain carrying empty pSET152. The full BGC makes thiocupinamide A (**1**) and thiocupinamide B (**2**). The production of both molecules is completely abolished and thiocupinamide C (**3**) is produced instead in the Δ*tcaO* mutant.

*S. coelicolor* M1146 carrying the complete *tca* BGC accumulated a major peak with *m/z* 1363.6040 (**1**) and a minor peak with *m/z* 1347.6085 (**2**) (**Figure 5C** and **Figure S6**). In contrast, the strain carrying pCAP03_P*tsrA*_*tca*_BGC_Δ*tcaO* accumulated a compound with *m/z* 1331.6137 (**3**, **Figure 5C**), as expected for a thioalbamide analogue with an isoleucine to valine substitution and a hydroxylation on the N,N-dimethyl histidinium (calc. *m/z* 1331.6131). The difference in masses compared to the products of the complete BGC (Δ31.9903 Da and Δ15.9955 Da, respectively) strongly suggests that the role of TcaO cupin is the sequential introduction of two oxygen atoms into the thiocupinamide structure. The Δ*tcaO* mutation could be genetically complemented with an intact copy of *tcaO* (**Figure S7**), supporting this notion. Additionally, MS/MS fragmentation data suggested that both oxygens are introduced on the first three residues of the core peptide (**Figures S8-10)**, although the absence of specific fragments covering that part of the compound hampered a more precise determination. Thus, we named these putative precursors thiocupinamide B (*m/z* 1347.6085) and thiocupinamide C (*m/z* 1331.6137). We could not detect the production of thiocupinamide A or any related thioamidated derivatives in the native *Streptomyces* sp. S.PNR29, which suggests that the *tca* BGC is silent in the assayed culture conditions and highlights the utility of heterologous BGC expression.

### Increasing thiocupinamide production by expressing multiple copies of the BGC

The lack of production in *Streptomyces* sp. S.PNR29 meant that direct comparisons with a native producer were not possible, although the heterologous production driven by P*tsrA* yielded much higher amounts of thiocupinamide in comparison to thioalbamide production by *A. alba*, as assessed by MS peak areas (**Figure 6**). To further increase the production levels, we explored the expression of additional copies of the refactored *tca* BGC. The instability of replicative plasmids in *Streptomyces* means they are not used for the expression of complex biosynthetic pathways, so stable expression of BGCs in *Streptomyces* requires integration of genes into specific genomic loci. For example, pCAP03 features φC31 integration elements that enable site-specific recombination with the *attB* integration site in *Streptomyces* genomes and carries a kanamycin resistance marker for selection. Therefore, to enable the expression of multiple copies of the BGC we created two new vectors (pSTW1 and pSTW2) compatible with pCAP03 and each other in terms of antibiotic resistance markers and integration sites but sharing the same overlaps for Gibson-like assembly (see Materials and Methods). Briefly, pSTW1 is based on pIJ10257^38^ (ϕBT1 integration and hygromycin resistance) and pSTW2 is based on pCMF92^39^ (ϕJoe integration and apramycin resistance). This allowed us to *de novo* assemble P*tsrA*_*tca_*BGC into pSTW1 and pSTW2 using the same PCR amplicons previously employed for the creation of pCAP03_P*tsrA*_*tca*_BGC. The resulting pSTW1_PtsrA_*tca*_BGC and pSTW2_P*tsrA*_*tca*_BGC constructs were combined with pCAP03_PtsrA_*tca*_BGC to create *S. coelicolor* M1146 strains carrying two or three refactored copies of the *tca* BGC. All these combinations boosted the production of thiocupinamide (**Figure 6**), in some cases up to 25-fold compared to thioalbamide production levels in *A. alba.* Notably, the inclusion of a third copy of the *tca* BGC did not increase the production respect to the combination of two copies, perhaps because of the pleiotropic effects associated to the integration of multiple plasmids in the *S. coelicolor* M1146 genome, or possibly due to a bottleneck associated with gene expression or metabolic flux.

**Figure 6.**
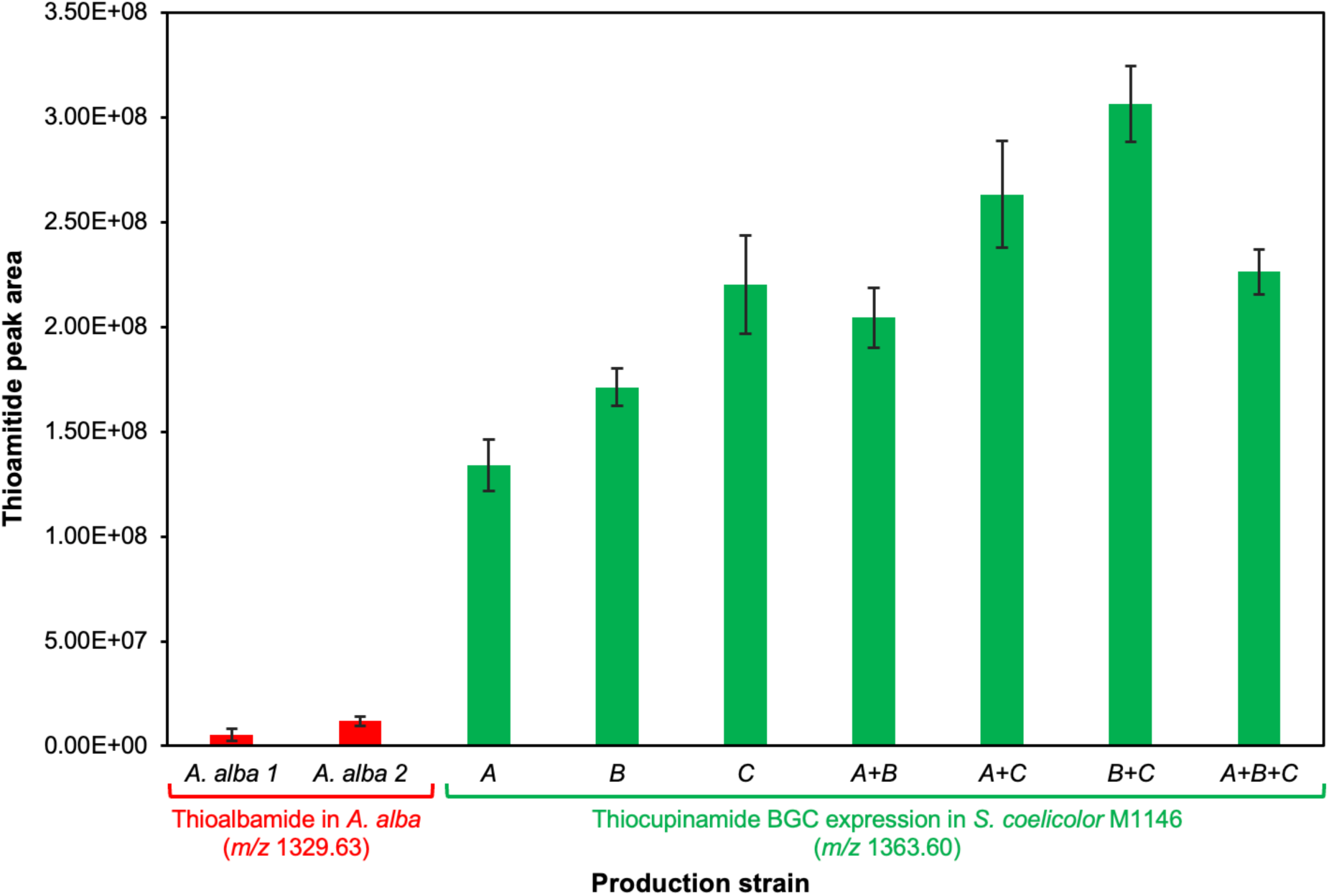
Multiple BGC copies increase thiocupinamide A production levels in *S. coelicolor* M1146. MS peak areas were compared to wild type thioalbamide production from two independent fermentations of *A. alba*. “*A. alba 1*” represents production from fermentations that were conducted in parallel to other fermentations in this experiment, while “*A. alba 2*” is the data from the fermentation shown in Figure 3. A = pCAP03 P*tsrA tca* BGC; B = pSTW1 P*tsrA tca* BGC; C = pSTW2 P*tsrA tca* BGC. Data from five fermentation replicates are shown, with vertical bars representing standard deviation.

### Thiocupinamide is a polyhydroxylated thioalbamide analogue

The high yields of thiocupinamide A produced by our engineered strains enabled the straightforward purification of a substantial amount of compound (11.3 mg) from a relatively small amount of SM12 solid medium (1.5 L) using a low-resolution debulking chromatography step followed by two HPLC steps (see Materials and Methods). A comprehensive set of NMR experiments (^1^H-NMR, DEPT-Q, HSQC, HMBC, and TOCSY, **Figures S11** to **S17**) was performed on 1.7 mg of pure thiocupinamide A. The analysis of these spectra revealed the connectivity expected for a thioalbamide-like molecule and allowed us to determine the modification introduced by TcaO as a two hydroxylations on the carbons β and γ1 of the Ile3 residue (**Figures 5B** and **7**, **Figures S18-S20**, **Table S5**). The isoleucine carbon β-hydroxylation is supported by DEPTQ, in which this carbon signal appears as a quaternary carbon (δ_C_ = 73.7, together with the absence of a correlation in HSQC) in contrast to the equivalent carbon in thioalbamide (δ_C_ = 36.8, appearing as a CH, δ_H_ = 2.13, m)^6^. The isoleucine carbon γ1 hydroxylation is supported by the change of its chemical shift in DEPTQ (δ_C_ = 72.9, appearing as a CH in HSQC, δ_H_ = 3.85, m) compared to the equivalent carbon in thioalbamide (δ_C_ = 26.6, appearing as a CH_2_ in HSQC, δ_H_ = 1.35, m and 1.64, m)^6^. HMBC and TOCSY correlations were fully consistent with the proposed structure (**Figure 7**, **Figures S18-S20**). The presence of a β-hydroxylation on isoleucine is consistent with an MS/MS fragment that results from a retro aldol fragmentation at this residue (**Figure S8**). An equivalent fragment is visible in the MS/MS spectrum of thiocupinamide B (**Figure S9**), along with a further retro aldol fragment from the β-hydroxylated histidinium (*m/z* 125.07). These data strongly suggest that thiocupinamide B lacks the hydroxylation on γ1 of isoleucine. To the best of our knowledge, isoleucine dihydroxylation is a very rare post-translational modification of ribosomal peptides that has only previously been observed in thiostrepton and related molecules like saalfelduracin, in which it is introduced by a cytochrome P450^40^ instead of a cupin-fold protein. The determination of the absolute stereochemistry of this modification in thiocupinamide will define if both modifications represent an example of convergent evolution or of biosynthetic stereoselective diversification. Absolute stereochemical determination of thioamitides has proven to be a long-standing challenge, due to the lack of crystal structures, as well as indefinite data from Marfey’s analyses^4,14^, which may result from the propensity of thioamides to racemise^41^. Further studies will be required to determine the true three-dimensional structures of all thioamitides, including thiocupinamide.

**Figure 7.**
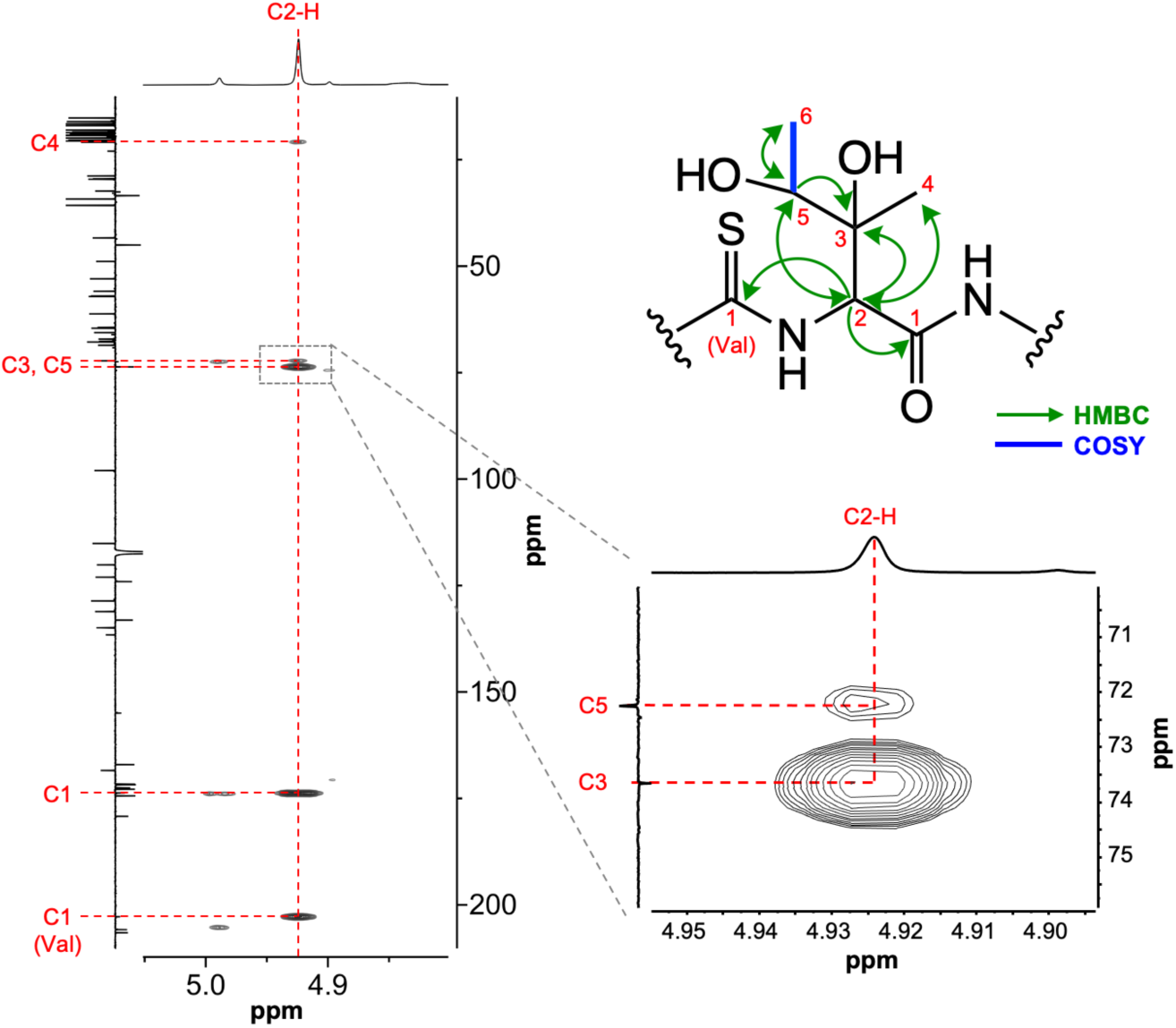
NMR characterisation of the dihydroxylated isoleucine residue. A section of the ^1^H-^13^C HMBC NMR spectrum is displayed showing the correlations of the α proton of this residue (C2-H). The full HMBC spectrum is shown in Figure S17 and 2D NMR correlations are shown in Figures S18-S20.

### Thiocupinamide A is a potent cytotoxic and antibacterial molecule

To investigate the biological activity of thiocupinamide A, we evaluated its potential to impair the viability of cancer cells. For this, the compound was tested across a broad panel of neoplastic cell lines, including highly aggressive and therapy-resistant tumours such as non-small cell lung cancer (NSCLC) (A549 and H460), glioblastoma multiforme (GBM) (U87-MG and T98G), and breast cancer (MDA-MB-231 and T47D). Specifically, cell viability was assessed using the MTT assay following 72 hours of exposure to increasing concentrations of thiocupinamide A (up to 10 µM). The results (**Figure 8A**) demonstrated a dose-dependent reduction in cell viability, with IC₅₀ values ranging from 5.1 to 54.5 nM, highlighting the potent cytotoxic activity of thiocupinamide A.

**Figure 8.**
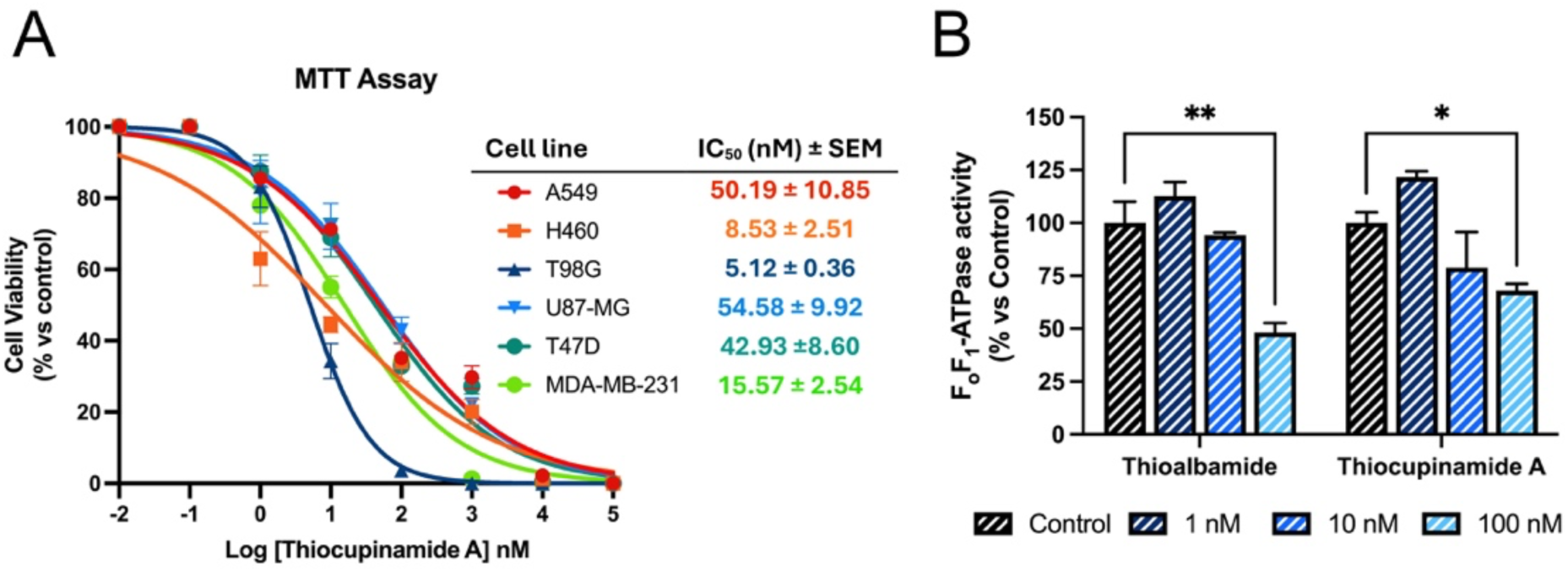
Biological activity of thiocupinamide A. (A) Cell viability assay (MTT) was performed on cancer cell lines exposed for 72 h to increasing concentrations of thiocupinamide A. IC_50_ ± SEM is reported for each cell line. (B) F_o_F_1_-ATPase activity in presence of different concentrations of thioalbamide or thiocupinamide A. Statistical significance was assessed by two-way ANOVA followed by Dunnett’s multiple comparisons test *, p<0.05, **, p<0.01.

Given that inhibition of oxidative phosphorylation by targeting the FoF₁-ATPase complex has been widely reported for this class of RiPP^7–9^, we next assessed the inhibitory potential of thiocupinamide A on the enzymatic activity of the mitochondrial FoF₁-ATPase *in vitro* (**Figure 8B**). The results revealed a marked inhibitory effect of this RiPP on this enzymatic complex, with its activity significantly reduced in the presence of 100 nM thiocupinamide A. Notably, this inhibitory activity was comparable to that observed for thioalbamide, as measured using the same spectrophotometric enzymatic assay, and aligns with the known mechanisms of action reported for other thioamitide compounds. The inhibition of F_o_F₁-ATPase and the resulting impairment of oxidative phosphorylation, responsible for mitochondrial ATP production in eukaryotic cells, may account for the differential sensitivity of the cancer cell lines to thiocupinamide A. It is well established that cancer cells undergo profound metabolic reprogramming, and that neoplastic diseases are characterized by substantial metabolic heterogeneity^42^. Although further investigation is needed, the varying sensitivity of the tested cancer cell lines to thiocupinamide may therefore reflect differences in their reliance on mitochondrial bioenergetic processes. It is noteworthy that the viability of the cell lines used turned out to be markedly reduced at concentrations lower than those required to inhibit the isolated enzyme. This phenomenon may be explained by the fact that partial inhibition of a key enzyme can disrupt essential metabolic or signalling networks, leading to cell death even in the absence of complete enzymatic inhibition. Additionally, potential off-target or cumulative effects, which cannot be excluded at this stage of the study, may further enhance cellular susceptibility.

The antibacterial activity of thiocupinamide A was evaluated against a panel of pathogenic Gram-positive bacteria (*Staphylococcus aureus* and *Streptococcus pyogenes*) and Gram-negative bacteria (*Pseudomonas aeruginosa* and *Klebsiella pneumoniae*), cultured in either Mueller-Hinton broth (MHB) or RPMI 1640 supplemented with 2% glucose (**Figure S21** and **Table 1**). Gram-negative strains were found to be resistant to thiocupinamide A at all tested concentrations (up to 64 µg/mL) in both culture media. In contrast, the RiPP exhibited activity against the Gram-positive strains, with minimum inhibitory concentration (MIC) values varying depending on the culture medium. Specifically, a MIC of 8 µg/mL was observed for both *S. aureus* and *S. pyogenes* in MHB, while a lower MIC of 1 µg/mL was recorded when the same strains were cultured in RPMI 1640 + 2% glucose. The results are consistent with the prior reports of the antimicrobial spectrum for other thioamitides^4,6^. Furthermore, these data indicate an enhanced antibacterial activity compared to that previously described for thioalbamide^6^, its closest structural analogue, suggesting how small structural differences can impact the biological activity of thioamitides.

**Table 1.**
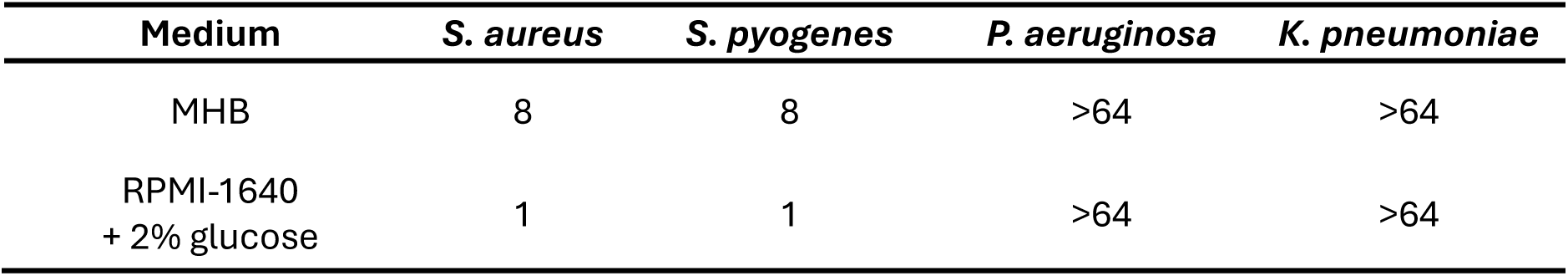
Antibacterial activity of thiocupinamide A. Data are presented as MIC values (µg/mL).

| Medium | *S. aureus* | *S. pyogenes* | *P. aeruginosa* | *K. pneumoniae* |
| --- | --- | --- | --- | --- |
| MHB | 8 | 8 | >64 | >64 |
| RPMI-1640<br>+ 2% glucose | 1 | 1 | >64 | >64 |

## CONCLUSION

The combination of quick *in vitro* refactoring and a curated selection of promoters has allowed us to overcome for the first time the challenging heterologous production of thioalbamide, as well as substantially increase its yield compared to the native producer. More generally, the infrequency of pathway-specific RiPP regulators and their compact BGCs makes RiPPs well suited to heterologous expression-based discovery and engineering. Our targeted search for new strong promoters has revealed P*tsrA* as an excellent tool for RiPP pathway refactoring. RiPP pathways are unusual amongst specialised metabolism as the precursor peptide is structural rather than catalytic, which necessitates high levels of gene expression. Along with these current results with P*tsrA*, we had previously shown that P*btmC* from the bottromycin BGC represented a highly active promoter^26^, yet RiPP pathway promoters remain an understudied resource for gene expression. One notable example is the use of the *godA* promoter from the goadsporin BGC^43^ for the expression of a lactazole BGC, which significantly enhanced pathway yield in heterologous expression^44^. We also showed that P*groEL2* represents a promising promoter for *Streptomyces* gene expression (**Figure 3**), which we had identified from *S. coelicolor* transcriptomic data^29^. It is worth noting that there is rarely a linear relationship between promoter activity and pathway yield, so it is useful to assess promoter libraries when expressing BGCs.

We have also shown how the simultaneous expression of two copies of the pathway in *Streptomyces* can also boost the production of thioamitides. Although the presence of multiple BGC copies is well documented among industrial overproducing strains generated by random mutagenesis^45,46^, it is underexplored as a rational approach in the metabolic engineering of *Streptomyces*, despite some successful examples, such as the use of *zouA*-dependent DNA amplification^46–48^ or the parallel expression of wild-type and refactored BGCs^49^. Here, we present pSTW1 and pSTW2, vectors sharing the highly efficient overhangs of pCAP03 for isothermal Gibson-like cloning, thus greatly facilitating the assembly and expression of up to three integrated copies of a given refactored BGC in *S. coelicolor*. The improvement we report in thioamitide yield is significant in the context of RiPP metabolic engineering^50^, especially considering the complexity of thioamitide biosynthesis and that it has been performed without any bioprocess optimisation in *S. coelicolor*. This result compares favourably to heterologous expression studies on darobactin in *Escherichia coli*^51,52^, where a 10-fold yield improvement for heterologous production compared to the native producer despite the highly optimised tools available for *E. coli*. Prior heterologous expression work on thioholgamide also provided a ~10-fold yield improvement^22^.

This heterologous expression platform can also be employed to accelerate the discovery of new thioamitides, as illustrated by the straightforward isolation of thiocupinamide A due to the high yields achieved by our system. The *tca* BGC initially attracted our attention due to the combination of a core precursor peptide almost identical to thioalbamide and the presence of tailoring enzymes suggesting the introduction of additional polar moieties on the final product of the pathway. Structural elucidation of thiocupinamide A corroborated that all four structural differences with thioalbamide (three additional hydroxylations and the lack of one methyl group due to the Ile/Val difference) increase the polarity of the molecule, making it more hydrophilic and therefore potentially more suitable as a formulable compound for drug development^53^. We showed that these modifications do not negatively impact cytotoxicity towards a variety of cancer cell lines (**Figure 8**), while antibacterial activity towards *Staphylococcus aureus* is enhanced ~3-fold compared to thioalbamide^6^.

It is important to identify new tailoring enzymes introducing hydrophilic modifications as part of pathways to naturally evolved thioamitides, as the introduction of charged residues via mutagenesis could potentially interfere with biological activity, and hydroxylated residues (Thr, Ser) are prone to heterocyclisation due to the promiscuity of the YcaO enzymes co-responsible for thioamidation^22^. Our identification of a pfam08007 hydroxylase highlights the versatility of cupin-fold oxygenases in oxidising unactivated carbon centres. Other members of this family include *E. coli* YcfD, which is a 2-oxoglutarate-dependent oxygenase that catalyses hydroxylation of an arginine residue of the ribosomal protein L16. Thus, the characterisation of new pathways constitutes a key step in the exploration of the structural diversity of these compounds towards their development as anticancer drugs in comparison to unnatural mutagenesis approaches, which are more likely to hamper the target binding. Our informatic analysis (**Figure 4**) indicates a rich diversity of tailoring enzymes and associated core peptides, so further work should aim to map the natural diversity of thioamitides and RiPPs from related pathways. These data would help to understand the structural determinants that are key for activity towards eukaryotic and prokaryotic targets.

## MATERIALS AND METHODS

### Chemicals

Unless otherwise specified, all chemical reagents were obtained from Sigma-Aldrich and all enzymes from New England Biolabs. Antibiotics were used at these final concentrations: 50 μg/mL kanamycin (Km), 50 μg/mL hygromycin (Hyg), 50 μg/mL carbenicillin, 50 μg/mL apramycin (Apra), 25 μg/mL chloramphenicol (Chl), and 25 μg/mL nalidixic acid. All primers and oligonucleotides were ordered desalted from IDT (Belgium). Ultrapure water was produced using a Milli-Q purification system (Merck) and all media and solutions were autoclaved prior to use, unless otherwise stated.

### Culture media

All ingredients were obtained from Sigma-Aldrich unless otherwise specified. Amounts are expressed in g/L unless otherwise specified and were prepared with ultrapure water.

**LB** (Lysogeny Broth): 10 tryptone, 5 yeast extract, 10 NaCl, pH 7. 15 agar was added for the solid version. **L** 10 tryptone, 5 yeast extract, 5 NaCl, 1 glucose, pH 7. **DNAm** (DifcoTM Nutrient Broth/Agar medium): purchased from Becton, Dickinson and Company. **SFM** (Soya Flour Mannitol): 20 soya flour (Holland and Barrett), 20 mannitol, 20 agar. **TSB** (Tryptic Soy Broth): Purchased from Oxoid. **SM12** (Screening Medium 12): 10 g soy flour, 50 g glucose, 4 g peptone, 4 g beef extract, 1 g yeast extract, 2.5 g NaCl, 5 g CaCO_3_, adjust to pH 7.6 with KOH. **YPDA**: 10 bacto yeast extract, 20 bacto peptone, 20 glucose monohydrate, 40 mg L^-1^ adenine hemisulfate. 20 agar were added for the solid version. **SD+CSM-Trp**: 1.7 YNB-AA-(NH_4_)_2_SO_4_ (Formedium), 5 (NH_4_)_2_SO_4_, 20 glucose, 20 agar, 20 mg L^-1^ adenine, and 740 mg L^-1^ CSM-Trp (Formedium).

### General molecular biology work

Unless otherwise specified, DNA fragments used for molecular cloning were always purified from agarose gels using the QIAquick™ gel extraction kit (Qiagen). Genomic DNA (gDNA) was extracted with a soil DNA extraction kit (FastDNA™ Spin Kit for soil, MP Biomedicals), and plasmid DNA was prepared using the Wizard® Plus SV Minipreps DNA Purification System (Promega). When required for cloning, PCR fragments were amplified using a high-fidelity DNA polymerase (Q5, New England Biolabs) and GoTaq DNA polymerase (Promega) was employed for analytical colony PCRs. All oligonucleotides are listed in **Table S1** and all plasmids are listed in **Table S2**.

### Bioinformatic identification of thioamitide BGCs

HopA1-like proteins from previously identified thioamitide BGCs (accessions: WP_020634200.1, TFI52254.1, WP_091117926.1, WP_017595620.1, WP_106236741.1, WP_155252669.1, WP_184984772.1, WP_190061747.1, WP_046417231.1, WP_167359510.1, BAN83919.1, KPC82023.1, WP_030193573.1, WP_018962004.1, WP_071375576.1, WP_039653089.1, WP_065960366.1) were each used as queries for BlastP^54^ searches using the NCBI non-redundant protein database with default search parameters in November 2024. The top 100 hits from each search were filtered for identity >30%, and accession lists were then combined and filtered for duplicates to provide 278 unique accessions. Protein sequences were obtained using ncbi-acc-download (https://github.com/kblin/ncbi-acc-download) and then filtered using a 97% sequence identity cut-off using CD-Hit^55^ to reduce redundancy. The resulting sequences were size-filtered to remove any proteins shorter than 200 amino acids to yield a dataset with 185 HopA1 accessions whose identity to each other was lower than 97%.

This accession list was analysed using RiPPER^33^ in Docker (https://hub.docker.com/r/streptomyces/rippertest) with default RiPPER parameters (minPPlen = 20, maxPPlen = 120, flanklen = 17.5, sameStrandReward = 5, maxDistFromTE = 8000, fastaOutputLimit = 3, prodigalScoreThresh = 7.5). This analysis provided 35 kb Genbank files centred on each accession and a list of putative precursor peptides for each BGC. Peptide sequences and associated information from the RiPPER output is provided as **Supplementary File 1**. The precursor peptide similarity network generated by RiPPER was visualised using Cytoscape^56^ (version 3.10.1) using the Prefuse Force Directed layout with HIT%ID. Predicted core peptides for each thioamitide subfamily were visualised using WebLogo^57^ (version 2.8.2) as a frequency plot. A custom colour scheme reflecting the residue physicochemical properties was generated based on features of the IMGT (International Immunogenetics Information System) and Zappo colour schemes (amino acids and hex colour codes in parentheses): aliphatic (AILV) = blue (#1B04AC), hydroxyl (ST) = green (#1B7837), sulfur (CM) = orange (#FFCC00), acidic (DE) = pale blue (#CCECFF), amide (NQ) = brown (#CCA504), basic (RHK) = red (#EC1504), aromatic (FWY) = purple (#CC99FF), conformationally special (GP) = pink (#FF00FF). To visualise the synteny and diversity of the associated BGCs, representative clusters were visualised using clinker^58^ (version 0.0.28).

### Strains

*S. coelicolor* M1146 was used as a heterologous expression host. *A. alba* DSM 44262, *Streptomyces varsoviensis* DSM 40346, *Streptomyces scabies* DSM 41658 and *S. laurentii* DSM 41684 were acquired from the German Collection of Microorganisms and Cell Cultures (DSMZ, Germany) and used as genetic source for the *taa* BGC, and the P*varA*, P*botA* and P*tsrA* promoters, respectively. *Streptomyces* sp. NRRL S4 was acquired from the ARS culture collection (NRRL, USA) and used as genetic source for the P*tsaA* promoter. *Streptomyces* sp. S.PNR29 was used as a genetic source for the *tca* BGC and was previously isolated from cultivated soil in Thailand^36^. Unless otherwise specified, all *Streptom*yces strain were grown in SFM (solid) and TSB (liquid) media at 28 °C. Spores and mycelium stocks were kept at −20 °C and −70 °C in 20% glycerol. *Saccharomyces cerevisiae* VL6–48N was used for transformation-associated recombination (TAR) cloning and was grown at 30 °C with shaking at 250 rpm in YPDA medium. Recombinant yeast selection was performed using selective media SD+CSM-Trp complemented with 5-fluoorotic acid (Fluorochem, 1 mg mL^−1^). Yeast cell stocks were kept at −70 °C in 20% glycerol. *E. coli* DH5α was used for standard DNA manipulations. *E. coli* ET12567/pR9604^59,60^ and *E. coli* ET12567/pUZ8002^61^ were used to transfer DNA to *Streptomyces* by intergeneric conjugation as previously described^62^. All *E. coli* strains were grown in LB medium at 37 °C. *E. coli* hygromycin selection was performed in DNAm (solid) and L (liquid) medium. *E. coli* cell stocks were kept at −20 °C and −70 °C in 20% glycerol.

### Pathway cloning and refactoring

From the curated selection of 10 promoters used for BGC refactoring, a total of six (P*ermE*\*, P*kasO*\*, P*groEL2*, P*hrdB*, PSF14 and P*accIV*) were ordered as synthetic DNA cassettes (Twist Biosciences, USA) with a general spacer-terminator-spacer-promoter-RBS-start codon structure (see **Table S3**), employing as spacers and terminators well-established synthetic biology elements^63^ previously tested in *Streptomyces* systems^64^. The other four promoters (P*botA*, P*tsrA*, P*tsaA* and P*varA*) were cloned from genomes of *S. scabies*, *S. laurentii*, *Streptomyces* sp. NRRL S4 and *S. varsoviensis*, respectively. Both the natural promoters and the synthetic cassettes were PCR-amplified (combining primers 1 and 2 for P*varA*, 3 and 4 for P*botA*, 5 and 6 for P*tsrA*, 7 and 8 for P*tsaA*, 9 and 10 for PSF14, 11 and 12 for P*ermE*\*, 13 and 14 for P*groEL2*, 15 and 16 for P*hrdB*, 17 and 18 for P*accIV* and 19 and 20 for P*kasO*\*, see **Table S1**).

The *taa* BGC was split into two PCR amplicons (of 4 and 6.5 kb) from *A. alba* gDNA using primers 21 and 22, and 23 and 24, respectively. For each refactored construct, these two fragments were assembled with the corresponding promoter-containing fragment and BamHI-EcoRI digested pSET152 using the HiFi assembly kit (NEB). Plasmids extracted from PCR-positive colonies (GoTaq, Promega) were checked by restriction analyses and then sequenced (Plasmidsaurus).

To create pCAP03_PtsrA_tca_BGC, the *tca* BGC was split into three PCR amplicons (ca. 4 kb each) using primers 27 and 28, 29 and 30, and 31 and 32, respectively (**Table S1**). For the generation of the Δ*tcaO* version of the pathway, the third *tca* BGC fragment was generated using primers 31 and 33 (**Table S1**). P*tsrA* was PCR-amplified from the genome of *S. laurentii* using primers 25 and 26 (**Table S1**). Both in the case of *tca* BGC and *tca* BGC Δ*tcaO*, four fragments were assembled into NdeI-XhoI digested pCAP03 using the HiFi assembly kit (NEB). Plasmids extracted from PCR-positive colonies (GoTaq, Promega) were checked by restriction analyses and then sequenced (Plasmidsaurus).

### Creation of pSTW1 and pSTW2

We had previously observed in the lab that the overhangs left by the NdeI – XhoI digestion of the trifunctional pCAP03 vector seem to be especially efficient for isothermal assemblies. To facilitate the assembly and refactoring of the thiocupinamide BGC into vectors compatible with pCAP03 (KmR, φC31 integration site), the region of pCAP03 containing these overhangs was cloned into pIJ10257 (HygR, φBT1 integration site) to create pSTW1, and into pCMF92 (ApraR, φJoe integration site) to create pSTW2.

To create pSTW1, a fragment of 1502 bp was amplified from pCAP03 by PCR using primers 36 and 37 (**Table S1**), digested using KpnI and HindIII, and cloned by ligation (T4 DNA ligase, Invitrogen) into pIJ10257 previously digested with the same restriction enzymes.

To create pSTW2, a fragment of 1502 bp was amplified from pCAP03 by PCR using primers 38 and 37 (**Table S1**), gel purified, digested using EcoRI and cloned by ligation into pCMF92 previously digested with EcoRI and PvuII. An undesirable XhoI additional restriction site situated within the apramycin resistance gene of pCMF92 was then removed by PCR mutagenesis using primers 39 and 40 (**Table S1**). XhoI and NdeI could therefore be used for plasmid linearization to generate the desired isothermal assembly cloning vector backbone.

Thus, to create pSTW1_P*tsrA*_*tca*_BGC and pSTW2_P*tsrA*_*tca*_BGC, the same PCR fragments employed to create pCAP03_P*tsrA*_*tca*_BGC (see previous section) were assembled into NdeI-XhoI digested pSTW1 and pSTW2 using the HiFi assembly kit (NEB).

### Generation of additional genetic constructs

To create pLF026, a PCR fragment was amplified from *A. alba* gDNA using primers 41 and 42, gel purified and cloned by Gibson-like isothermal assembly (NEBuilder HiFi DNA assembly kit, New England Biolabs) into NdeI-HindIII digested pIJ10257, which was then gel purified and confirmed by sequencing (Plasmidsaurus). To create pIJ10257_*tca*O, a PCR fragment was amplified from *Streptomyces* sp. S.PNR29 using primers 34 and 35, gel purified, digested with NdeI and HindII, and cloned by ligation (T4 DNA ligase, Invitrogen) into NdeI-HindIII digested pIJ10257, which was then gel purified and confirmed by sequencing (Plasmidsaurus).

To create pCAP03_*taa*_BGC, a fragment of the *A. alba* genome was captured into pCAP03 employing TAR cloning, following the same protocol described previously for other thioamitide BGCs^6^. Briefly, a vector to capture the desired genomic DNA digestion fragment was constructed by the use of a modified (ligase-free) Gibson assembly between a XhoI-NdeI linearized pCAP03 and oligonucleotides 43 and 44. The resulting capture vector was linearized between the capture arms using PmeI and was then co-transformed after gel purification into *S. cerevisiae* VL6-48N spheroplasts together with *A. alba* gDNA digested using MluI. Successful BGC capture by pCAP03 was confirmed by colony PCR and then by restriction analysis and sequencing (Plasmidsaurus) of the extracted construct (pCAP03_*taa*_BGC). *E. coli* ET12567/pR9604 was transformed with pCAP03_*taa*_BGC by electroporation, and transformants were then used to transfer pCAP03_*taa*_BGC into *S. coelicolor* M1146 by intergeneric conjugation^62^. Apramycin-resistant *S. coelicolor* M1146 exconjugants containing integrated pCAP03_*taa*_BGC were verified by PCR.

### Semi-automated *Streptomyces* conjugation

*E. coli* ET12567/pUZ8002 was transformed with the ten thioalbamide BGC constructs (each carrying different promoters) alongside pSET152 as positive transformation/conjugation control using an automated bacterial transformation protocol performed on the CyBio Felix platform. Briefly, 5 μL of competent cells were transformed with plasmid DNA, recovered with 90 μL of SOC followed by plating of 7 μL cells in 12-well CytoOne® Multiwell plates containing LB + antibiotics (Apra 50 μg/mL, Km 50 μg/mL, Chl 25 μg/mL). Colonies were manually picked and grown in 1.5 mL LB + antibiotics (Apra 50 μg/mL, Km 50 μg/mL, Chl 25 μg/mL) in a 96-deep well plate. Overnight cultures were used to generate glycerol stocks. An Opentrons OT-2 protocol was designed to perform the automated conjugation (Conjugation_protocol_simplified_Thioalbamidev3.json; **Supplementary File 2**) as follows: the donor *E. coli* strains were grown overnight in 1.5 mL of LB + antibiotics in a 96-deepwell plate then 50 μL of culture were transferred to a 96-well microtiter plate (4titude-0-116, 300 µL round well format), spun down for 3 minutes at max speed, washed with 100 μL LB twice before being resuspended in 50 μL LB. For each reaction, 10 μL of *S. coelicolor* M1146 spores were transferred to a new plate, heat shocked at 50 °C for 10 minutes before 20 μL of *E. coli* were added to them. Spores and *E. coli* were spun down for 3 min at 5,000 x *g* (manual step) then resuspended in 50 μL that were plated in 6-well CytoOne® Multiwell SFM plates supplemented with 10 mM MgCl_2_ and 50 µg/mL cycloheximide with the Hamilton Microlab STAR liquid handling platform. Antibiotic overlay was performed with the Pontons OT2 platform by adding antibiotics in the 6-well CytoOne® Multiwell plates. The plates were incubated at 28 °C. *Streptomyces* exconjugants grew for all the reactions performed. The exconjugants obtained were manually streaked in petri dishes containing SFM media supplemented with Apra (50 μg/mL) and nalidixic acid (25 μg/mL), then incubated at 28 °C. Colony picking was performed manually, 4 colonies for each conjugation were inoculated in TSB + antibiotics in 24-well plates then incubated at 30 °C with shaking at 200 rpm. For the strains with pSET152 the antibiotics were Apra (50 μg/mL) and nalidixic acid (25 μg/mL) while for those with the pCAP03 were Km (50 μg/mL) and nalidixic acid (25 μg/mL).

### Fermentation, extraction and detection of thioamitides

*A. alba* DSM 44262, *Streptomyces* sp. S.PNR29 and *S. coelicolor* M1146 strains carrying the refactored *taa* and *tca* BGCs were grown in SFM solid medium for sporulation. After harvesting spores using established methods^62^, spores were spread on SM12 solid medium (33 mL of medium in a 90 mm petri dish) for thioamitide production. Production cultures were grown for 10 days at 30 °C. For analytical scale thioamitide extraction, one plug of agar was extracted employing the wide end of a sterile blue micropipette tip, placed in a 2 mL Eppendorf tube and mixed with 1 mL of methanol. These samples were vigorously shaken for one hour using a vortex platform and centrifuged at 14,000 rpm for 10 minutes in a microcentrifuge. 500 µL of supernatant were transferred into a 1.5 mL HPLC vial for LC-MS analysis.

LC-MS analysis was performed using an Agilent 1290 Infinity II HPLC coupled to an Agilent 6546 LC/QTOF. Samples (5 µL) were injected onto a Phenomenex Luna Omega polar C18 1.6 µm (50 x 2.1 mm) column set at 40 °C. The HPLC method was based on linear gradient of 0-95% of acetonitrile in water + 0.1% formic acid over 6 minutes (flow rate 0.6 mL/min). Ionization was carried out using the Dual AJS/ESI mode and positive detection mode was employed. Mass spectrometry data was collected between *m/z* 50 and 1700, and data-dependent MS/MS data was collected using collision-induced dissociation (CID). The MS parameters were as follows: gas temperature = 320 °C, drying gas = 8 L/min, nebulizer = 35 psi, sheath gas temperature = 350 °C, sheath gas flow = 11 L/min, capillary voltage = 3500 V, nozzle voltage = 1000 V, fragmentor voltage = 210 V, skimmer voltage = 100 V, octopole RF voltage = 750 V, collision cell gas = N_2_ (22 psi). Stepped CID settings between 35 and 100 were assessed for optimal MS/MS fragmentation.

### Purification and structural elucidation of thiocupinamide A

Fresh spores of *S. coelicolor* M1146 carrying pSTW2_P*tsrA*_*tca*_BGC were spread on 1.5 L of SM12 solid medium (split on six 30 x 30 cm square plates containing 250 mL each) and grown at 30 °C for 10 days. After this, the agar was cut into small pieces with a scalpel, transferred into a bottle and extracted twice with 1.5 L of methanol, shaking vigorously for 30 minutes. After centrifugation (30 minutes, 20,000 x *g*), the methanolic supernatant was dried *in vacuo*, and the solid residue redissolved in 100 mL of 50% (v/v) methanol/water. This concentrated crude extract was centrifuged (30 min, 8,000 rpm) and the supernatant was loaded into a low-resolution chromatography column (Biotage, Sfär C18 D Duo 100 Å 30 µm, 120 g) for debulking using a 0.1% formic acid in water (solvent A) /0.1% formic acid in acetonitrile (solvent B) gradient from 5 to 100 % B over 10 column volumes. After analysis by LC-MS (see method above), fractions (45 mL each) containing thiocupinamide A were dried in a Genevac evaporator, and combined by dissolution in 5 mL of 50% (v/v) methanol/water. Thiocupinamide A was purified from this solution by two HPLC steps using a Kinetex XB-C18 5 µm 100 Å (250 x 10 mm) column in an Agilent 1290 infinity II HPLC system. In both HPLC purification steps thiocupinamide A was automatically collected following DAD detection at 272 nm. In the first HPLC step, a 0.1% formic acid in water (A) / acetonitrile (B) elution gradient was used (T = 0 min, 5% B; T = 2 min, 5% B; T = 6 min, 40% B; T = 25, 98% B; T = 28, 5% B. Thiocupinamide A retention time: 12.1 min). In the second HPLC step, a 0.1% formic acid in water (A) / methanol (B) elution gradient was used (T = 0 min, 5% B; T = 2 min, 5% B; T = 6 min, 70% B; T = 18, 70% B; T = 20, 95% B; T = 22, 95% B; T = 22.10, 5% B. Thiocupinamide A retention time: 12.6 min). The collected material was freeze dried to afford 11.3 mg of pure thiocupinamide A in the form of a white powder.

### NMR data acquisition

For NMR data acquisition, 1.7 mg of thiocupinamide A was dissolved in 200 µL of acetonitrile-d3 and transferred into a 2 mm NMR tube. NMR experiments were conducted on a Bruker Avance Neo 600 MHz spectrometer equipped with a TCI cryoprobe at 298 K. Spectra were analysed using Mnova version 16 (Mestrelab research). The NMR experiments conducted were ^1^H-NMR (16 scans), ^13^C-NMR (18,000 scans), DEPT-Q (18,000 scans) TOCSY (10 scans), HSQC (10 scans) and HMBC (24 scans).

### Cell cultures

All the cell lines used in this work were purchased from the American Culture Collection (ATCC, Manassas, VA). For maintenance purposes, glioblastoma cells (U87-MG and T98G) were cultured in DMEM high-glucose, lung cancer cells (A549 and H460) in RPM I1640, and breast cancer cells (MDA-MB-231 and T47D) in DMEM-F12. All the media were supplemented with 10% fetal bovine serum, 2 mM L-glutamine and 1% penicillin/streptomycin. Media and supplements were all from Merck KGaA, Darmstadt, Germany. Treatments were performed in the above-mentioned media containing a lower amount of supplemented serum (2%). All cell lines were cultured at 37 °C in 5% CO_2_ in a humidified atmosphere.

### Cell viability assay

Cell viability was assessed using the MTT (3-(4,5-dimethylthiazol-2-yl)-2,5-diphenyltetrazolium bromide) assay. Cells were seeded in 48-well plates at a density of 2×10⁴ cells/well and incubated overnight. They were then treated with varying concentrations of thiocupinamide A for 72 hours, with DMSO as vehicle control. At the end of the treatment, MTT (0.5 mg/mL) was added to each well, and plates were incubated at 37 °C for 2 h to allow formazan crystal formation. Crystals were solubilized in DMSO, and absorbance was measured at 570 nm using a microplate reader. Dose-response curves were generated by non-linear regression (GraphPad Prism 9) to calculate IC₅₀ values.

### F_o_F_1_-ATPase activity assay

The ability of the tested compound to inhibit the activity of the F_o_F_1_-ATPase complex was tested using the MitoTox™ Complex V OXPHOS Activity Assay Kit (Abcam, ab109907), following the manufacturer’s instructions.

### Determination of MIC values

The minimum inhibitory concentration (MIC) of tested compound was determined by the CLSI (Clinical and Laboratory Standards Institute) broth microdilution method. Briefly, overnight cultures of *S. aureus* ATCC 25923*, S. pyogenes* ATCC 19615, *K. pneumonia* ATCC 13883, and *P. aeruginosa* ATCC 27853 were diluted to an optical density at 600 nm of 0.1 and were added in a 96-well plate containing serial dilutions of the compound. The test was performed both in Muller Hinton (MH) broth and in RPMI 1640 supplemented with 2% glucose. Bacterial growth was determined visually after overnight incubation at 37 °C.

## ACKNOWLEDGEMENTS

This work was supported by a Biotechnology and Biological Sciences Research Council (BBSRC) Institute Strategic Programme Grant *Harnessing Biosynthesis for Sustainable Food and Health* (HBio; BB/X01097X/1) awarded to the John Innes Centre (JIC), a BBSRC research grant (BB/V016024/1) awarded to A.W.T., and a BBSRC National Bioscience Research Infrastructure Grant (BBS/E/ER/23NB0007) awarded to the Earlham Institute. Further financial support was provided by the National Recovery and Resilience Plan (PNRR), Mission 4 “Education and Research” – Component 2 “From Research to Enterprise”, Investment 1.1, through the PRIN-PNRR 2022 project (Project code #P2022BLSMY), awarded to A.R.C. We are very grateful for the technical assistance at JIC provided by Lionel Hill for LC-MS, Sergey Nepogodiev for NMR, Govind Chandra for informatics and Hairuo Li for input into the genome mining. We thank Vladimir Larionov (National Cancer Institute, NIH, USA) for S. cerevisiae VL6-48N and Bradley Moore (Scripps Institution of Oceanography, University of California San Diego, USA) for pCAP03.

