## Supplementary Information for "Efficient *in vitro* refactoring and biosynthetic gene cluster amplification for the overproduction and accelerated discovery of anticancer thioamitides"

### SUPPLEMENTARY TABLES

**Table S1** Oligonucleotides used in this study.

| Number[a] | Sequence (5' to 3')[b] | Application |
|---|---|---|
| 1 | AAACAGCTATGACATGATTACGAATTCGAT**CGATGGGCTCTCCTCTGGGGCTTCG** | Cloning of P*varA* for Gibson assembly |
| 2 | CGGTGCTACGGCATCGTCATGCGAATTCAT**GAGGGCCTTTTCCTTTCCTTACGGGTTTGG** | |
| 3 | AAACAGCTATGACATGATTACGAATTCGAT**GCGACCTCTCGCCTCTCTGCGCACAG** | Cloning of P*botC* for Gibson assembly |
| 4 | CGGTGCTACGGCATCGTCATGCGAATTCAT**GGTGTGGTCCTTTCGTCTGTCTTCTCGTCTG** | |
| 5 | AAACAGCTATGACATGATTACGAATTCGAT**GGGTTGGCTTCGCCACGCTCAGG** | Cloning of P*tsrA* for Gibson assembly |
| 6 | CGGTGCTACGGCATCGTCATGCGAATTCAT**GCCGTTCACAACCTTTCGGCAAGTC** | |
| 7 | AAACAGCTATGACATGATTACGAATTCGAT**CTGTGGAAGCCTCAGGACTCACTCG** | Cloning of P*tsaA* for Gibson assembly |
| 8 | CGGTGCTACGGCATCGTCATGCGAATTCAT**GCTTCAACCTTCCCGCTAATCGAATC** | |
| 9 | AAACAGCTATGACATGATTACGAATTCGAT**TGCAGTTTTATTCTCTCGCCAGCACTG** | Amplification of PSF14 from synthetic cassette for Gibson assembly |
| 10 | CGGTGCTACGGCATCGTCATGCGAATTCAT**ATGGGGCCTCCTGTTCTAGACGATCC** | |
| 11 | AAACAGCTATGACATGATTACGAATTCGAT**CAAACACCCCATGTCGATACTGAACG** | Amplification of P*ermE** from synthetic cassette for Gibson assembly |
| 12 | CGGTGCTACGGCATCGTCATGCGAATTCAT**ATGGGGCCTCCTGTTCTAGACGATCC** | |
| 13 | AAACAGCTATGACATGATTACGAATTCGAT**CCATTGTAATAGCCACCAAAAGAGTGATG** | Amplification of P*groEL* from synthetic cassette for Gibson assembly |
| 14 | CGGTGCTACGGCATCGTCATGCGAATTCAT**ATGCATGTGAAGTGGTCCCTCCAGG** | |
| 15 | AAACAGCTATGACATGATTACGAATTCGAT**TGTCACGCTAGGAGGCAATTCTATAAG** | Amplification of P*hrdB* from synthetic cassette for Gibson assembly |
| 16 | CGGTGCTACGGCATCGTCATGCGAATTCAT**GAACAACCTCTCGGAACGTTGAAAAACGG** | |
| 17 | AAACAGCTATGACATGATTACGAATTCGAT**GCGACTTCCAGAGAAGAAGACTACTG** | Amplification of P*accIV* from synthetic cassette for Gibson assembly |
| 18 | CGGTGCTACGGCATCGTCATGCGAATTCAT**GTGTAGTCCTCCTTCGTATTGCACGAC** | |
| 19 | AAACAGCTATGACATGATTACGAATTCGAT**ACGCCACGCGTAGTGAGACATACAC** | Amplification of P*kasO* from synthetic cassette for Gibson assembly |
| 20 | CGGTGCTACGGCATCGTCATGCGAATTCAT**ATGGTCACCTCTTCAACTCAGATACTAG** | |
| 21 | **ATGAATTCGCATGACGATGCCGTAGCACCG** | Cloning of amplicon 1 of *taa* BGC |
| 22 | **CGCCCGCTGGATGTGCTTCATGTGGGCGGGCACGTCCTCGTC** | |
| 23 | **GACGAGGACGTGCCCGCCCACATGAAGCACATCCAGCGGGCG** | Cloning of amplicon 2 of *taa* BGC |
| 24 | CCAAGCTTGGGCTGCAGGTCGACTCTAGAG**CCGAAGTGGACATACGAGGCGGTGGAC** | |
| 25 | TTTGACGCCTCCCATGGTATAAATAGTGGC**GGGTTGGCTTCGCCACGCTCAGG** | |

| | | |
|---|---|---|
| 26 | CACCACGTCTTCTTCGTTCTTCTCCAT**GCCGTTCACAACCTTTCGGCAAGTC** | Cloning of P*tsrA* for Gibson assembly with *tca* BGC |
| 27 | GACTTGCCGAAAGGTTGTGAACGGC**ATGGAGAAGAACGAAGAAGACGTGGTG** | Cloning of amplicon 1 of *tca* BGC |
| 28 | CCGTGTTCGCGCCGTCCAGTTGGAC**GATCAGATCGGCTCTCCTCGTCAGCACC** | |
| 29 | GGTGCTGACGAGGAGAGCCGATCTGATC**GTCCAACTGGACGGCGCGAACACGG** | Cloning of amplicon 2 of *tca* BGC |
| 30 | GGACGAAGTCGCCTACCACCTGGGAG**GTGACGTAGTTGAACAACACCGAGGG** | |
| 31 | CCCTCGGTGTTGTTCAACTACGTCAC**CTCCCAGGTGGTAGGCGACTTCGTCC** | Cloning of amplicon 3 of *tca* BGC |
| 32 | GCACGTTCCTTATATGTAGCTTTCGACATA**CAGGTTACCCCGTTGGATCGCC** | |
| 33 | GCACGTTCCTTATATGTAGCTTTCGACATA**GGGACTCCCGCTTTCGTTCGG** | Cloning of amplicon 3 of *tca* BGC without *tcaO* |
| 34 | GATACAGATACACAT**ATGTACGTCCCCGAGTTCACCGACC** | Cloning *tcaO* in pIJ10257 |
| 35 | GATACAGATACAAGCTT**CCCTTCTGCGCCTCAACCGGC** | |
| 36 | GATACAGATACAGGTACC**CGCAAAAACTCGGTTTGACGCCTCCC** | Amplification of pCAP03 region for generation of pSTW1 and/or pSTW2 |
| 37 | GATACAGATACAAAGCTT**GACTAGGATGAGTAGCAGCACGTTCC** | |
| 38 | GATACAGATACAGAATTC**CGCAAAAACTCGGTTTGACGCCTCCC** | |
| 39 | **GGTACGCGTCGATTATCT**G**GAGAATGACCACTGCTGTGAGCGCTTTGC** | Removal of XhoI restriction site in pCMF92 |
| 40 | **CTCACAGCAGTGGTCATTCTC**C**AGATAATCGACGCGTACCAACTTGCCATCC** | |
| 41 | GGTAGGATCGTCTAGAACAGGACACCCATATGAGATACGAGGCCCCATAT**GAGATACGAGGTTCTTGGTTCGCTG** | Amplification of regulator adjacent to *taa* BGC |
| 42 | GCTCATGAGAACCCTAGGGGATCCAAGCTT**TCAGGAGGCCAGGCGGTCC** | |
| 43 | TTTGACGCCTCCCATGGTATAAATAGTGGCTCGAGGCGCTGGTGGTGATCACCGGTGCCGCCAACGTCACCGGCGAAGTGTGGCCGTTTAAACGTGACTCTCGC | Construction of pCAP03 capture vector for TAR cloning of *taa* BGC |
| 44 | AGCAGCACGTTCCTTATATGTAGCTTTCGACATATGGCCTCGCGTGAGCGAAGGGCGCCACAGACTCCCAGGACCGCGAGAGTCACGTTTAAACGGCCACACTTC | |

a. Numbers are referred to in the methods section.
b. Primer annealing regions are highlighted in bold and restriction sites are underlined.

**Table S2** Plasmids used in this study.

| Plasmid | Features | Application | Resistance Marker |
|---|---|---|---|
| pCAP03[1] | ΦC31 integration in *Streptomyces*, replicative in *E. coli* and in yeast | Cloning shuttle vector | Kanamycin |
| pCAP03_*taa*_BGC | *taa* BGC under the control of its native promoter | Expression of *taa* BGC | Kanamycin |
| pCAP03_P*tsrA*_*tca*_BGC | *tca* BGC under the control of the P*tsrA* promoter | Expression of *tca* BGC | Kanamycin |
| pCAP03_P*tsrA*_*tca*_BGC_Δ*tcaO* | truncated *tca* BGC under the control of thr P*tsrA* promoter | Expression of a truncated *tca* BGC lacking *tcaO* | Kanamycin |
| pIJ10257[2] | ΦBT1 integration in *Streptomyces*, replicative in *E. coli,* includes P*ermE** | Cloning shuttle vector | Hygromycin |
| pIJ10257_*tcaO* | *tcaO* under the control of P*ermE** | Complementation of the Δ*tcaO* mutant | Hygromycin |
| pLF026 | putative regulator adjacent to the *taa* BGC under the control of P*ermE** within the pIJ10257 backbone | Expression of the putative *taa* BGC-adjacent regulator | Hygromycin |
| pR9604[3] | *tra* genes. *E. coli* replication | Conjugation helper plasmid | Carbenicillin |
| pSET152[4] | ΦC31 integration in *Streptomyces*, replicative in *E. coli* | Cloning shuttle vector | Apramycin |
| pSET152_ P*accIV* _*taa*_BGC | *taa* BGC under P*accIV* | Expression of *taa* BGC | Apramycin |
| pSET152_ P*botA*_*taa*_BGC | *taa* BGC under P*botA* | Expression of *taa* BGC | Apramycin |
| pSET152_ P*ermE**_*taa*_BGC | *taa* BGC under P*ermE** | Expression of *taa* BGC | Apramycin |
| pSET152_ P*hrdB*_*taa*_BGC | *taa* BGC under P*hrdB* | Expression of *taa* BGC | Apramycin |
| pSET152_ P*groEL2*_*taa*_BGC | *taa* BGC under P*groEL2* | Expression of *taa* BGC | Apramycin |
| pSET152_ P*kasO**_*taa*_BGC | *taa* BGC under P*kasO** | Expression of *taa* BGC | Apramycin |
| pSET152_ Psf14_*taa*_BGC | *taa* BGC under Psf14 | Expression of *taa* BGC | Apramycin |
| pSET152_ P*tsaA*_*taa*_BGC | *taa* BGC under P*tsaA* | Expression of *taa* BGC | Apramycin |
| pSET152_ P*tsrA*_*taa*_BGC | *taa* BGC under P*tsrA* | Expression of *taa* BGC | Apramycin |
| pSET152_ P*varA*_*taa*_BGC | *taa* BGC under P*varA* | Expression of *taa* BGC | Apramycin |
| pSTW1 | ΦBT1 integration in *Streptomyces*, replicative in *E. coli* | Cloning shuttle vector | Hygromycin |
| pSTW1_P*tsrA*_*tca*_BGC | tca BGC under P*tsrA* | Expression of *tca* BGC | Hygromycin |
| pSTW2 | ΦJoe integration in *Streptomyces*, replicative in *E. coli* | Cloning shuttle vector | Apramycin |
| pSTW2_P*tsrA*_*tca*_BGC | *tca* BGC under P*tsrA* | Expression of *tca* BGC | Apramycin |
| pUZ8002[5] | *tra* genes. *E. coli* replication | Conjugation helper plasmid | Kanamycin |

**Table S3** Synthetic promoter cassettes used for BGC expression.

| Promoter | Construction | Sequence[a] |
|---|---|---|
| PSF14 | Cassette_spacer/Ba_0015/spacer/PSF14 | GATACAGAATTCTGCAGTTTTATTCTCTCGCCAGCACTGTAATAGGCACTAACCAGGCATCAAATAAAACGAAAGGCTCAGTCGAAAGACTGGGCCTTTCGTTTTATCTGTTGTTTGTCGGTGAACGCTCTCTACTAGAGTCACACTGGCTCACCTTCGGGTGGGCCTTTCTGCGTTTATAGCGTGCGTACACCTTAATCACCGCTTCATGCTAAGGTCCTGGCTGCATGCGCGGTCGATCTTGACGGCTGGCGAGAGGTGCGGGGAGGATCTGACCGACGCGGTCCACACGTGGCACCGCGATGCTGTTGTGGGCACAATCGTGCCGGTTGGTAGGATCGTCTAGAAC**AGGAGG**CCCCATATGGATACA |
| *PermE** | Cassette_spacer/L3S2P21/spacer/PermE* | GATACAGAATTCCAAACACCCCATGTCGATACTGAACGAATCGACGCACACTCCCTTCCTTGCTCGGTACCAAATTCCAGAAAAGAGGCCTCCCGAAAGGGGGGCCTTTTTTCGTTTTGGTCCCATTTTTGCCTTGCGACAGACCTCCTACTTAGATTGCCACGCGGTCGATCTTGACGGCTGGCGAGAGGTGCGGGGAGGATCTGACCGACGCGGTCCACACGTGGCACCGCGATGCTGTTGTGGGCACAATCGTGCCGGTTGGTAGGATCGTCTAGAAC**AGGAGG**CCCCATATGGATACA |
| P*hrdB* | Casette_spacer/L1U1H09/spacer/PhrdB | GATACAGAATTCTGTCACGCTAGGAGGCAATTCTATAAGAATGCACACTGCACGACGATGTTCGCATCGTCGTTTTTTTTTCCGTGGTTGAGTCAGCGTCGAGCACGCGGCCCGCCTTCCGCCGGAACGGCGGGGTCCGGGCACGCCAAACCCCTCCTGTGGCTGTGGCCGGCCACCGCCGTCACCTTCGGACCCCGTGGAGCCGCTCCCGGTTCCACGGGGTCCGAAGGTGTGATGAGCAGGCTGCGCCTTCCTCGCGCGGCCGCAAGGTACGAGTTGATGACCTTGTTTATCCGCATCTGACCAATTTTGATCGCTTACGGGGTGTGACTCGGGCCACGCGGATTGGGCGTAACGCTCTTGGGAACAACACGATGACCTAAGAGGTGACAGCCGCGGAGGGAATACGGACGCCGTTCACGGCGCTGTGCATCTCCCCGGCCCGCCCGCACCGTCGGCCCATTCCCAAGCCGGTGGTCGGCCCCTGTCCGCCGTGGACGGGGCCGGAAGCCGTTTTTCAACGTTCC**GAGAGG**TTGTTCATGCATATGGATACA |
| P*groEL2* | Casette_spacer/L3S1P11/spacer/PgroEL2 | GATACAGAATTCCGCGACTTCCAGAGAAGAAGACTACTGACTTGAGCGTTCCGGTCTCATCGAGACGAACAATAAGGCCTCCCAAATCGGGGGGCCTTTTTATTTTTCAACAAAATGTTAGAGACCACTACGAGATTTGAGGTAAACCAAATAAGCACGTAGTGGCCACCACCGACTATTTGCAACAGTGCCGTTGATCGTGCTATGATCGACTGATGTCATCAGCGGTGGAGTGCAATGTCGTGCAATACG**AAGGAGG**ACTACACATGCATATGGATACA |
| P*kasO** | Casette_spacer/T7t/spacer/PkasO* | GATACAGAATTCACGCCACGCGTAGTGAGACATACACGTTCGTTGGGTTCACTACTCTAACCCCATCGGCCGTCTTAGGGGTTTTTTGTTCGAGAAACAAGGCAGTTCCGGGCTGAAAGTAGCGCCGGGGTCGACTCTAGAGCTGAGTTGGCTGCTGCCACCGCTGAGCAATAACTAGCATAACCCCTTGGGGCCTCTAAACGGGTCTTGAGGGGTTTTTTGCTGAAAGGAGGAACTATATCCGCGGGATCCTGTTCACATTCGAACGGTCTCTGCTTTGACAACATGCTGTGCGGTGTTGTAAAGTCGTGGCCAGGAGAATACGACAGCGTGCAGGACTGGGGGAGTTACTAGTATCTGAGTT**GAAGAGG**TGACCATATGGATACA |
| P*accIV* | Casette_spacer/L3S1P32/spacer/PaccIV | GATACAGAATTCCGCGACTTCCAGAGAAGAAGACTACTGACTTGAGCGTTCCGGTCTCATCGAGACGAACAATAAGGCCTCCCAAATCGGGGGGCCTTTTTATTTTTCAACAAAATGTTAGAGACCACTACGAGATTTGAGGTAAACCAAATAAGCACGTAGTGGCCACCACCGACTATTTGCAACAGTGCCGTTGATCGTGCTATGATCGACTGATGTCATCAGCGGTGGAGTGCAATGTCGTGCAATACG**AAGGAGG**ACTACACATGCATATGGATACA |

a. All features shown in these synthetic sequences were retained during the cloning process apart from the nucleotides that followed the start codons, which were replaced with the first gene of the cloned BGC.

**Table S4** Predicted functions of the proteins encoded in the *Streptomyces* sp. S.PNR29 thiocupinamide BGC.

| Gene name | Protein accession number | Pfam/NCBI domain[a] | Predicted biosynthetic function | Size (AA) |
|---|---|---|---|---|
| *tcaA* | WP_289934973.1 | NF033415 | Thioamitide-family precursor peptide | 84 |
| *tcaC* | WP_289934974.1 | PF01636 | Phosphotransferase catalysing Ser/Thr dehydration with TcaD | 367 |
| *tcaD* | WP_289934975.1 | PF17914 | HopA1-like lyase catalysing Ser/Thr dehydration with TcaC | 328 |
| *tcaE* | WP_289934976.1 | CD05154 | Phosphotransferase-like cyclase | 295 |
| *tcaF* | WP_289934977.1 | PF02441 | Flavoprotein catalysing Cys oxidative decarboxylation. | 198 |
| *tcaG* | WP_289934978.1 | PF06325 | Methyltransferase catalysing dimethylation of His12 | 409 |
| *tcaH* | WP_289934979.1 | PF02624 | YcaO protein catalysing thioamidation with TcaI | 450 |
| *tcaI* | WP_289934980.1 | PF07812 | TfuA protein catalysing thioamidation with TcaH | 225 |
| *tcaJ* | WP_289934981.1 | PF05721 | Oxygenase catalysing His12 hydroxylation | 279 |
| *tcaK* | WP_289934982.1 | PF00112 | Protease catalysing leader peptide removal | 250 |
| *tcaRed* | WP_289934983.1 | PF00106 | NADPH-dependent reductase catalysing pyruvyl reduction | 287 |
| *tcaCYP* | WP_289934984.1 | PF00067 | Cytochrome P450 catalysing Phe5 hydroxylation | 395 |
| *tcaO* | WP_289934985.1 | PF08007 | Cupin-fold oxygenase catalysing Ile3 dihydroxylation | 298 |

a. The top Pfam domain is listed based on NCBI CD-Search[6]. If no Pfam domain was detected with an E-value cutoff of 0.01, the top scoring domain from an alternative domain database is listed.

**Table S5** NMR assignments for thiocupinamide A in acetonitrile-d3. Figure S11 shows details of residue naming and atom numbering.

| Residue | $\delta_C$, mult. | $\delta_H$, mult. (*J*, in Hz) | Residue | $\delta_C$, mult. | $\delta_H$, mult. (*J*, in Hz) |
|---|---|---|---|---|---|
| **LA** | | | **avMCys** | | |
| 1 | 179.21, C | | 1 | 171.80, C | |
| 2 | 67.81, CH | 4.35 (q, 6.9) | 2 | 55.93, CH | 3.38 (m)[ol] |
| 3 | 19.93, $CH_3$ | 1.40 (d, 6.9) | 3 | 43.37, CH | 3.38 (m)[ol] |
| **Val 1** | | | 4 | 19.00, $CH_3$ | 1.16 (d, 5.8) |
| 1 | 202.76, C | | 5 | 98.01, CH | 5.44 (d, 7.0) |
| 2 | 65.60, CH | 4.78 (dd, 3.9, 8.9) | 6 | 134.92, CH | 7.29 (dd, 7.0, 10.6) |
| 3 | 32.36, CH | 2.85 (m) | NH (1) | | 8.22 (d, 5.9) |
| 4[a] | 19.33, $CH_3$ | 1.00 (d, 6.9) | NH (2) | | 10.69 (d, 10.6) |
| 5[a] | 15.18, $CH_3$ | 0.89 (d, 7.0) | **Val 3** | | |
| NH | | 7.88 (d, 9.0) | 1 | 172,86, C | |
| **Ile-2OH** | | | 2 | 64.57, CH | 3.86 (m)[ol] |
| 1 | 173.72, C | | 3 | 29.58, CH | 2.26, (m)[ol] |
| 2 | 67.14, CH | 4.92(s) | 4[a] | 20.62, $CH_3$ | 1.19 (d, 6.44) |
| 3 | 73.66, C | | 5[a] | 18.67, $CH_3$ | 1.02 (m)[ol] |
| 4 | 20.87, $CH_3$ | 1.10 (s) | NH | | 7.09 (d, 5.4) |
| 5 | 72.85, CH | 3.85 (m)[ol] | **Ala 2** | | |
| 6 | 17.01, $CH_3$ | 1.12 (d, 6.4) | 1 | 174.38, C | |
| NH | | ND | 2 | 52.96, CH | 4.00 (qd, 7.3, 4.6) |
| **Gly** | | | 3 | 16.57, $CH_3$ | 1.50 (d, 7.3) |
| 1 | 172.54, C | | NH | | 7.29 (m)[ol] |
| 2 | 45.05, $CH_2$ | 3.58 (d, 16.2) | **Val 4** | | |
| | | 3.74 (d, 16.2) | 1 | 171.67, C | |
| NH | | ND | 2 | 57.01, CH | 4.12 (dd, 9.2, 4.5) |
| **Phe-OH** | | | 3 | 28.80, CH | 2.26 (m)[ol] |
| 1 | 205.83, C | | 4[a] | 18.04, $CH_3$ | 0.78 (d, 7.0) |
| 2 | 67.64, CH | 4.66 (dd, 11.6, 2.9) | 5[a] | 16.05, $CH_3$ | 0.70 (d, 7.1) |
| 3 | 33.45, $CH_2$ | 3.00 (dd, 13.5, 2.9) | NH | | 7.48 (d, 9.2) |
| | | 3.38 (m)[ol] | **dmHis-OH** | | |
| 4 | 124.09, C | | 1 | 167.05, C | |
| 5 | 154.96, C | | 2 | 57.14, CH | 4.05 (dd, 7.6, 9.4) |
| 6 | 115.13, CH | 6.93 (m) | 3 | 61.22, CH | 5.98 (d, 9.4) |
| 7 | 128.61, CH | 7.14 (m) | 4 | 133.13, C | |
| 8 | 120.15, CH | 6.83 (m) | 5 | 123.03, CH | 7.67 (d, 1.7) |
| 9 | 131.14, CH | 7.18 (m) | 6 | 136.57, CH | 8.30 (d, 1.7) |
| NH | | | N1-Me | 34.25, $CH_3$ | 3.90(s) |
| **Ala 1** | | | N2-Me | 35.72, $CH_3$ | 3.89 (s) |
| 1 | 206.57, C | | NH | | 7.59 (d, 7.6) |
| 2 | 64.32, CH | 5.47 (q, 7.3) | | | |
| 3 | 18.51, $CH_3$ | 1.84 (d, 7.3) | | | |
| NH | | ND | | | |
| **Val 2** | | | | | |
| 1 | 171.78, C | | | | |
| 2 | 68.58, CH | 4.25 (d, 10.9) | | | |
| 3 | 29.50, CH | 2.26, (m)[ol] | | | |
| 4[a] | 19.20, $CH_3$ | 0.95 (d, 6.9) | | | |
| 5[a] | 20.40, $CH_3$ | 1.04 (m)[ol] | | | |
| NH | | ND | | | |

a. For each valine, these signals may be interchanged.
Abbreviations: ol = overlapped signal, ND = NH proton not detected due to deuterium exchange
Residues and noncanonical amino acids: LA = lactic acid, Ile-2OH = dihydroxyisoleucine, Phe-OH = o-hydroxy-phenylalanine, avMCys = 2-aminovinyl-3-methyl-cysteine; dmHis-OH = β-hydroxy-dimethyl histidinium

### SUPPLEMENTARY FIGURES

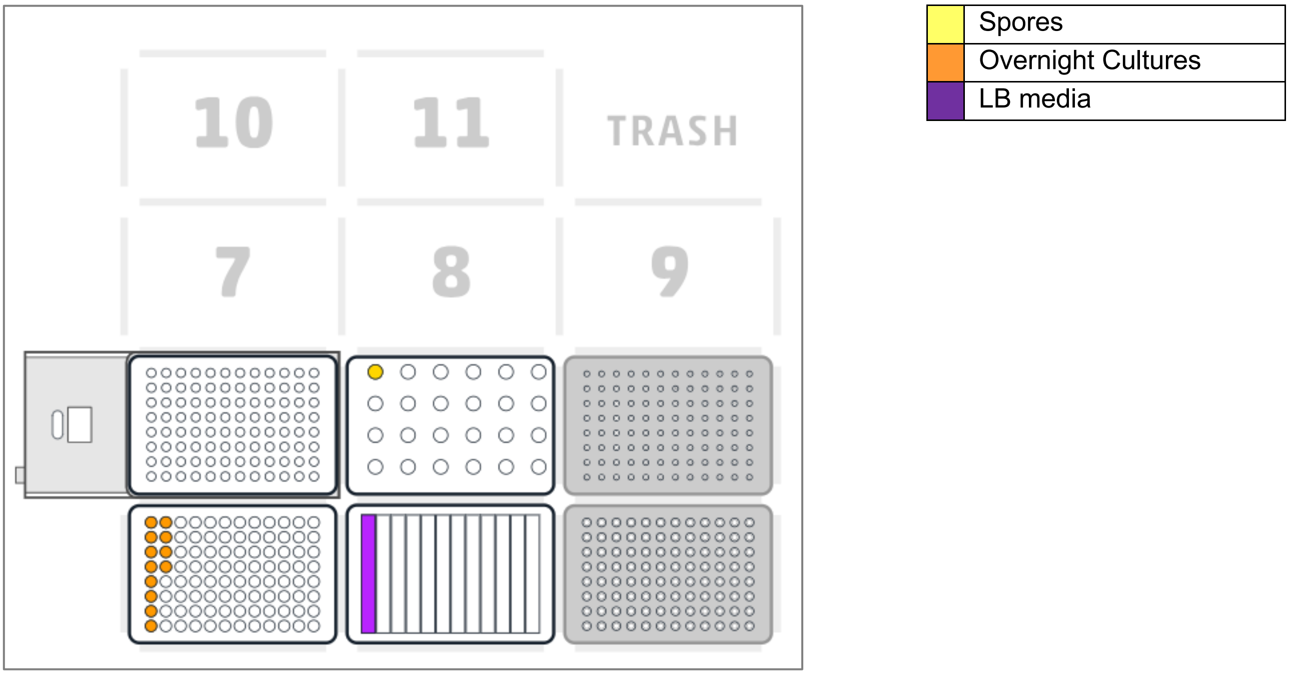


**Figure S1** Deck layout of the Opentrons OT-2 for the automated conjugation protocol. Position 1 contains overnight cultures with the donor strain. Position 2 holds a 12-well reservoir with LB in well A1. Position 4 contains the temperature block. Position 5 includes spores in an Eppendorf tube (well A1). Positions 3 and 8 contain the required Opentrons tips. The protocol is provided as **Supplementary File 2**.

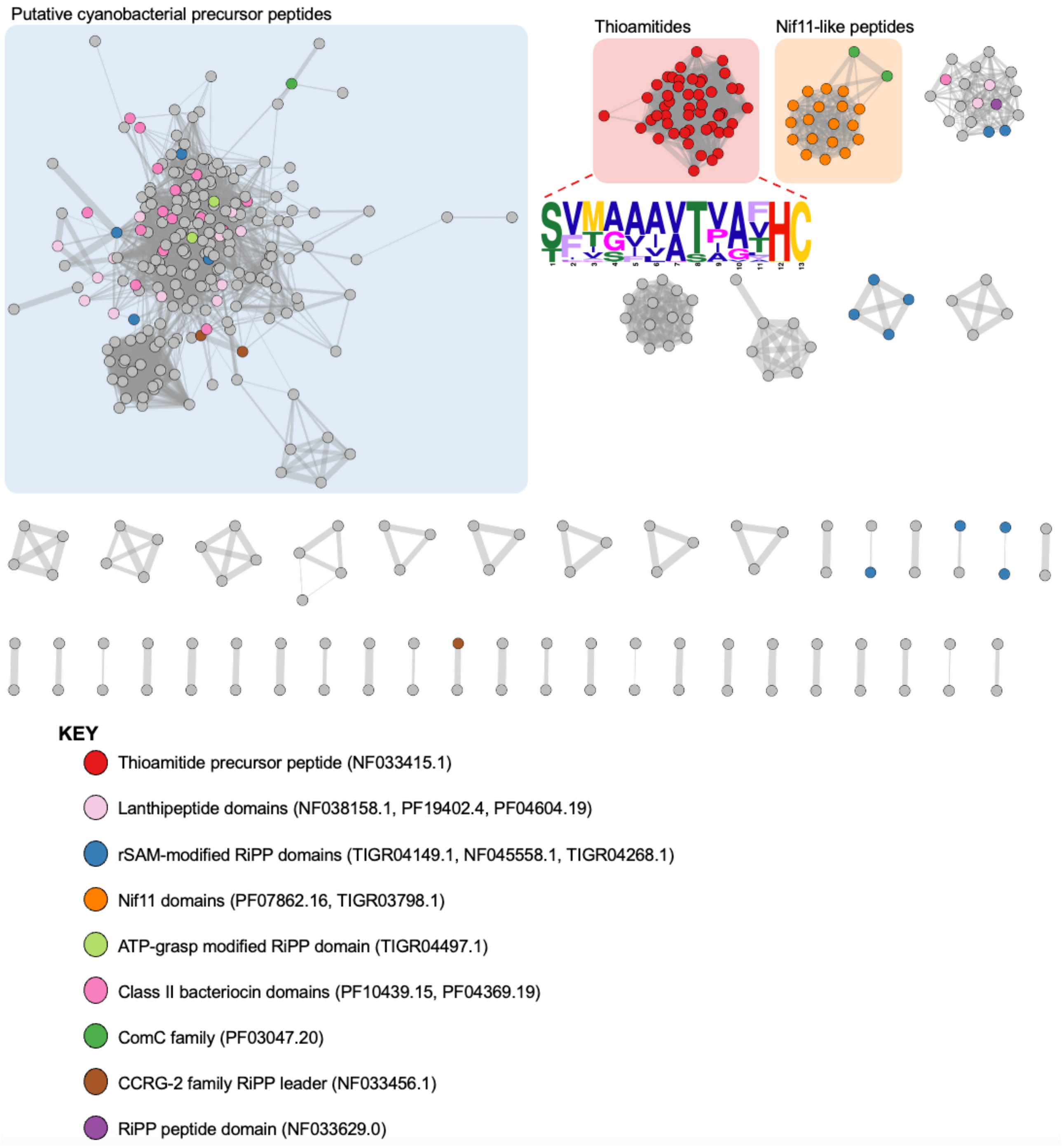


**Figure S2** Total precursor peptide network from RiPPER analysis[7] visualised using Cytoscape[8]. Each node is a short peptide and an edge is shown where there is at least 40% sequence identity between peptides. Edge thickness is scaled from 0.7 for %ID = 40 to 17.5 for %ID = 100. Nodes are colour coded if a putative RiPP precursor domain was identified and selected precursor peptide families are highlighted. We previously reported on the identification of largest network, which features cyanobacterial peptides. The sequence logo of all thioamitides with a predicted 13 amino acid core peptide is also shown (visualised using WebLogo[9]). The RiPPER output and associated network data are provided as **Supplementary File 1**.

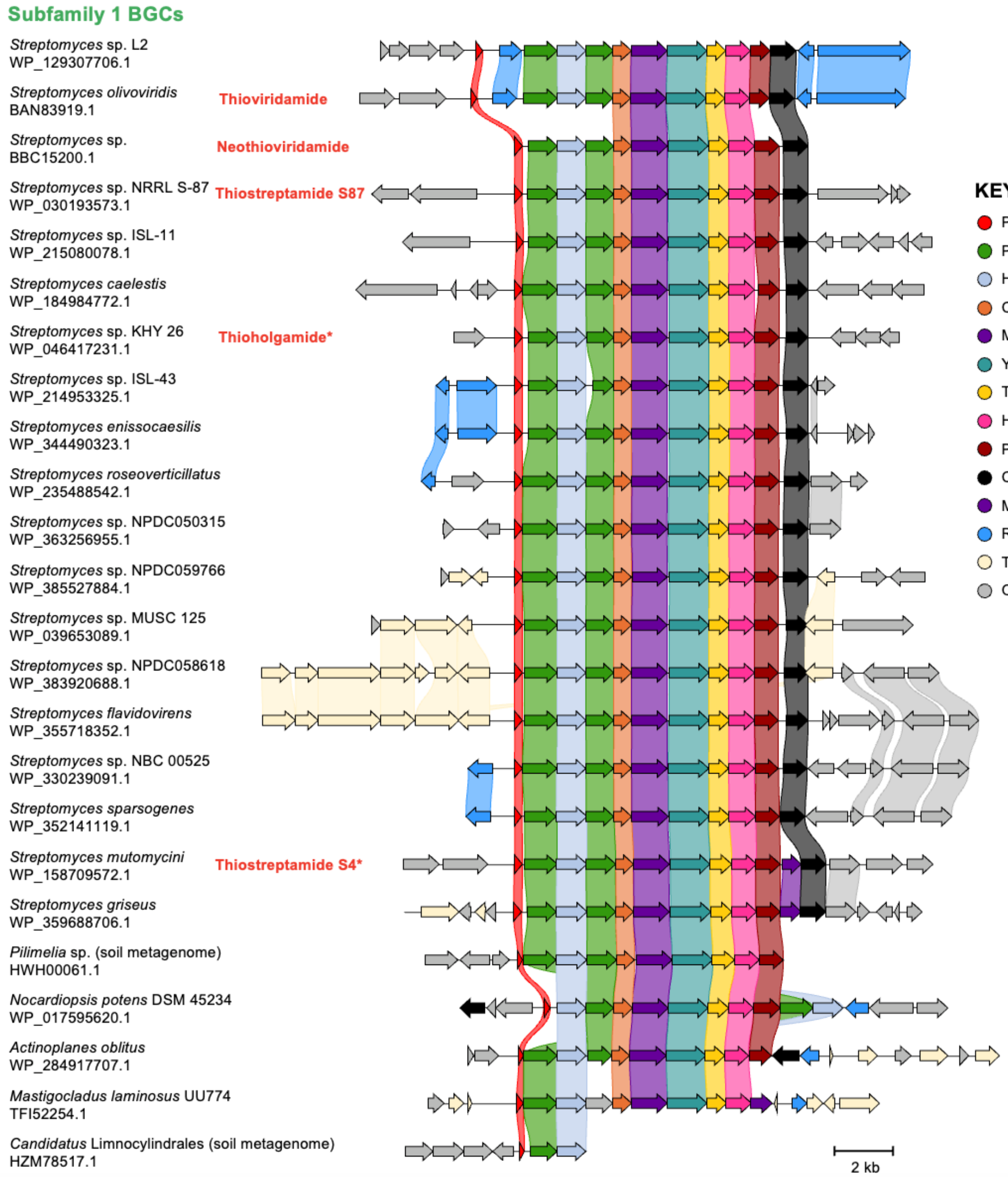


**Figure S3** Synteny of the BGCs from thioamitide subfamily 1 visualised using clinker[10]. Gene links are shown if identity is over 30% between encoded proteins. The accession of the BGC-encoded HopA1 protein is listed under each strain name. Selected flanking genes are shown. BGCs associated with known thioamitides are labelled, where an asterisk indicates that the HopA1 protein and precursor peptide are identical to the characterised BGC, but a BGC from a different organism is displayed.

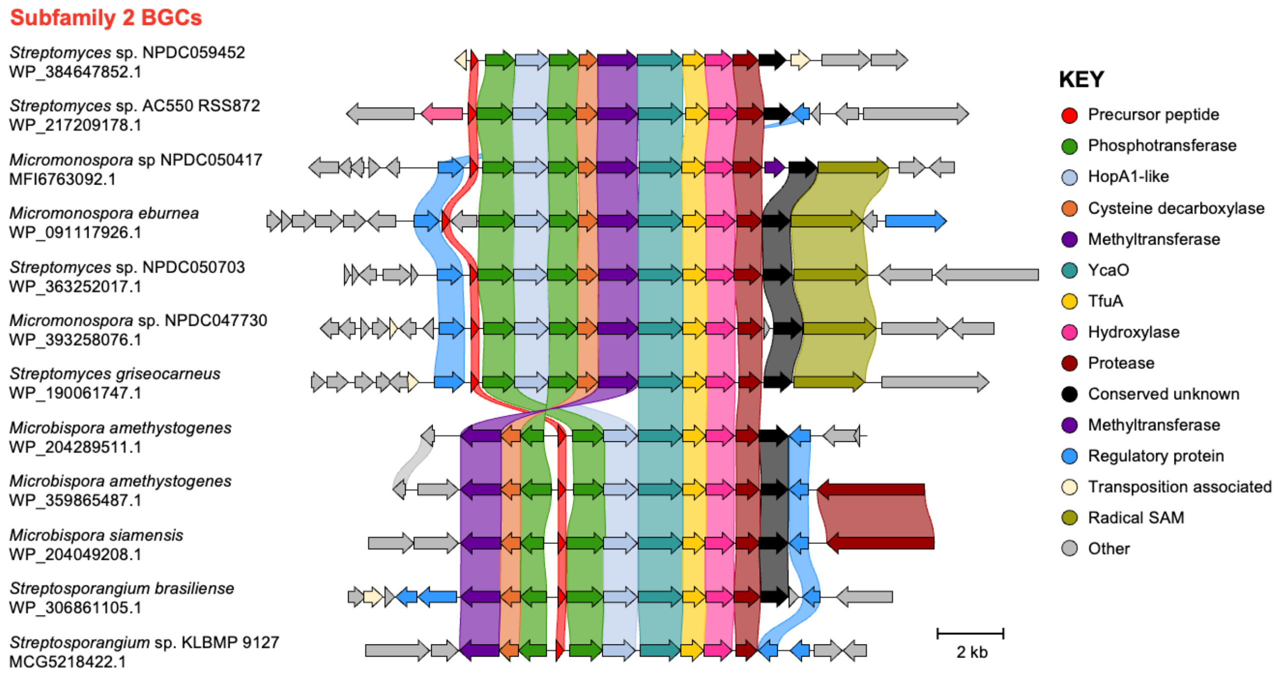


**Figure S4** Synteny of the BGCs from thioamitide subfamily 2 visualised using clinker[10]. The accession of the BGC-encoded HopA1 protein is listed under each strain name. Gene links are shown if identity is over 30% between encoded proteins. Selected flanking genes are shown.

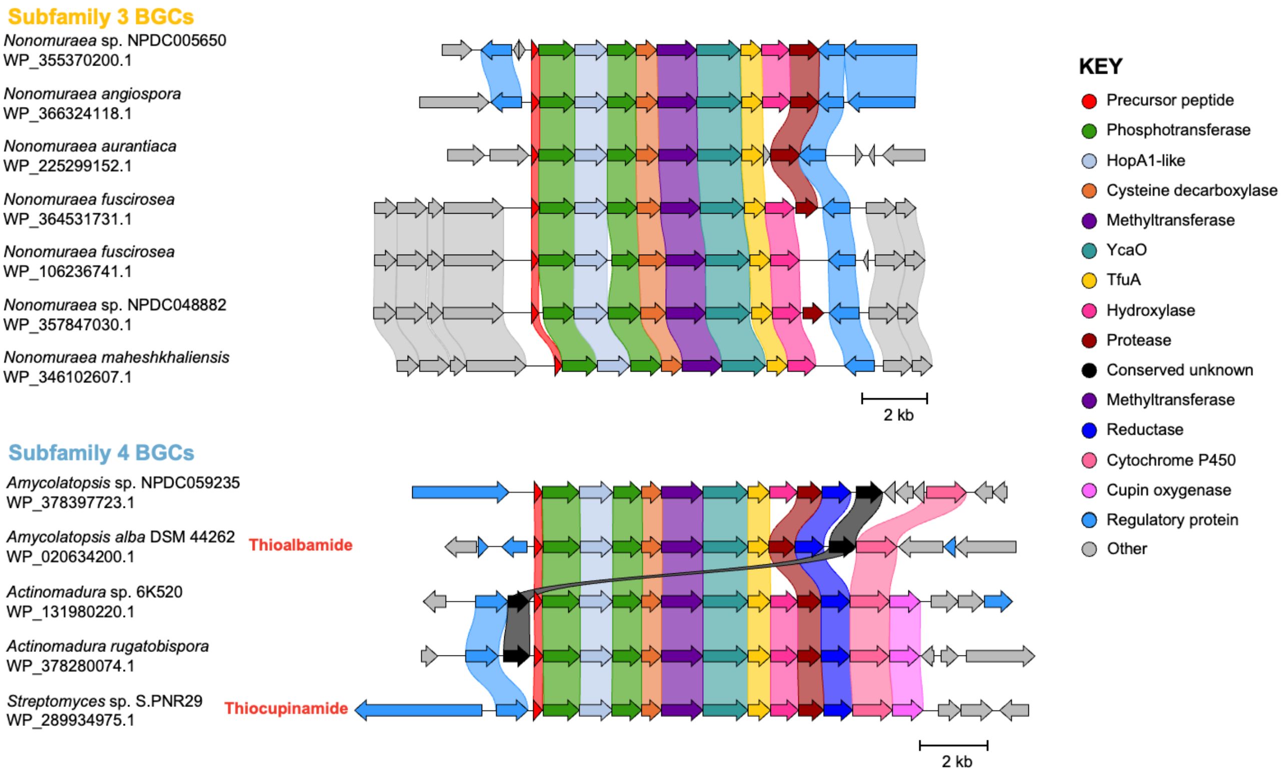


**Figure S5** Synteny of the BGCs from thioamitide subfamilies 3 and 4 visualised using clinker[10]. The accession of the BGC-encoded HopA1 protein is listed under each strain name. Gene links are shown if identity is over 30% between encoded proteins. Selected flanking genes are shown. BGCs associated with known thioamitides are labelled.

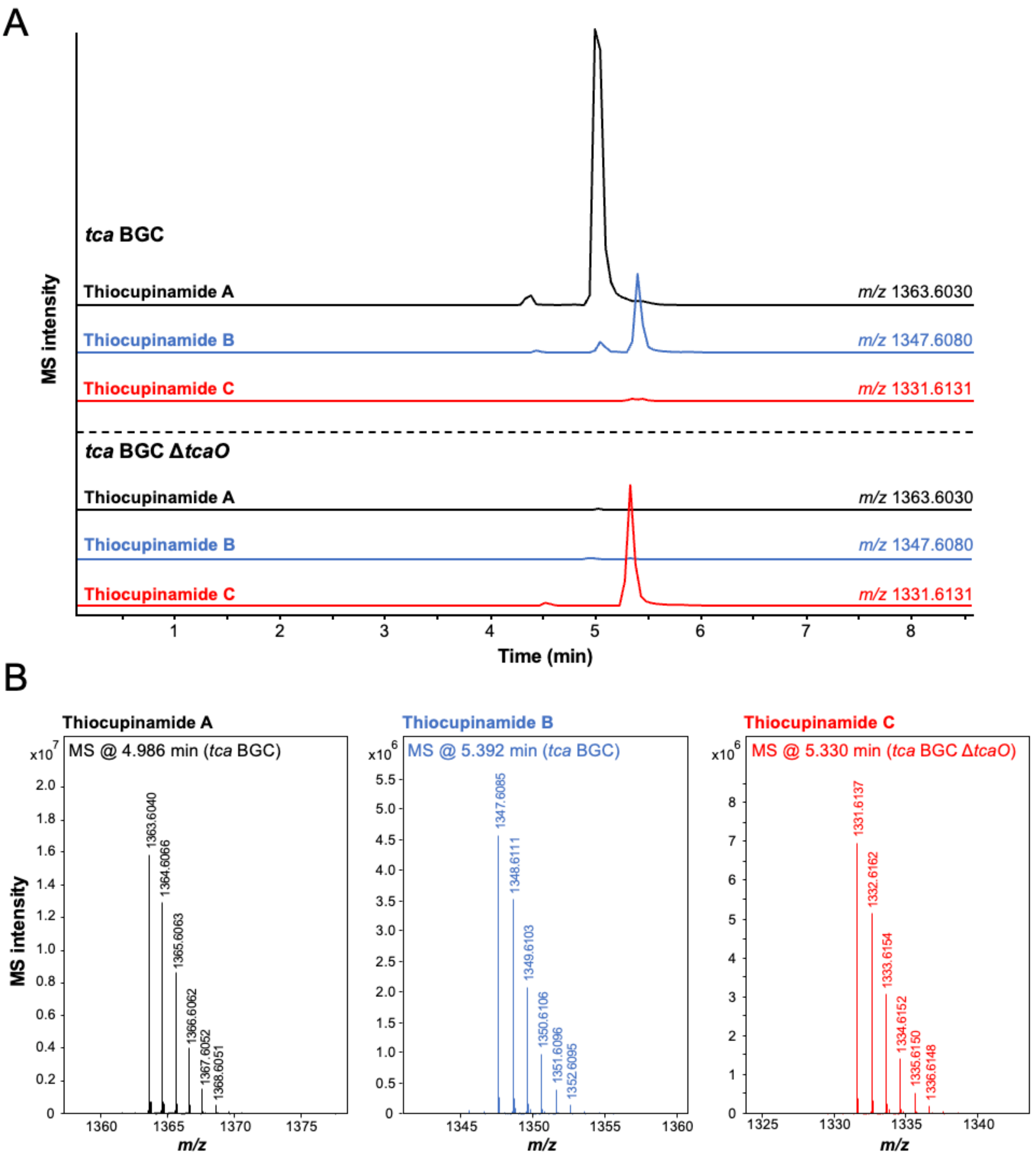


**Figure S6** Heterologous production of thiocupinamides. **A.** Extracted ion chromatograms showing the relative proportion of the molecules detected in *S. coelicolor* M1146 integrated with either pSET152 P*tsrA tca* BGC or pSET152 P*tsrA tca* BGC Δ*tcaO*. Extracted ion chromatograms are shown for each *m/z* (10 ppm extraction window) and are scaled relative to the largest overall peak. **B.** MS spectra for each molecule. Thiocupinamide A and B spectra are from the tca BGC expression experiment and the thiocupinamide C spectrum is from *tca* BGC Δ*tcaO*. Each spectrum is obtained from the apex of the respective peak shown in panel A.

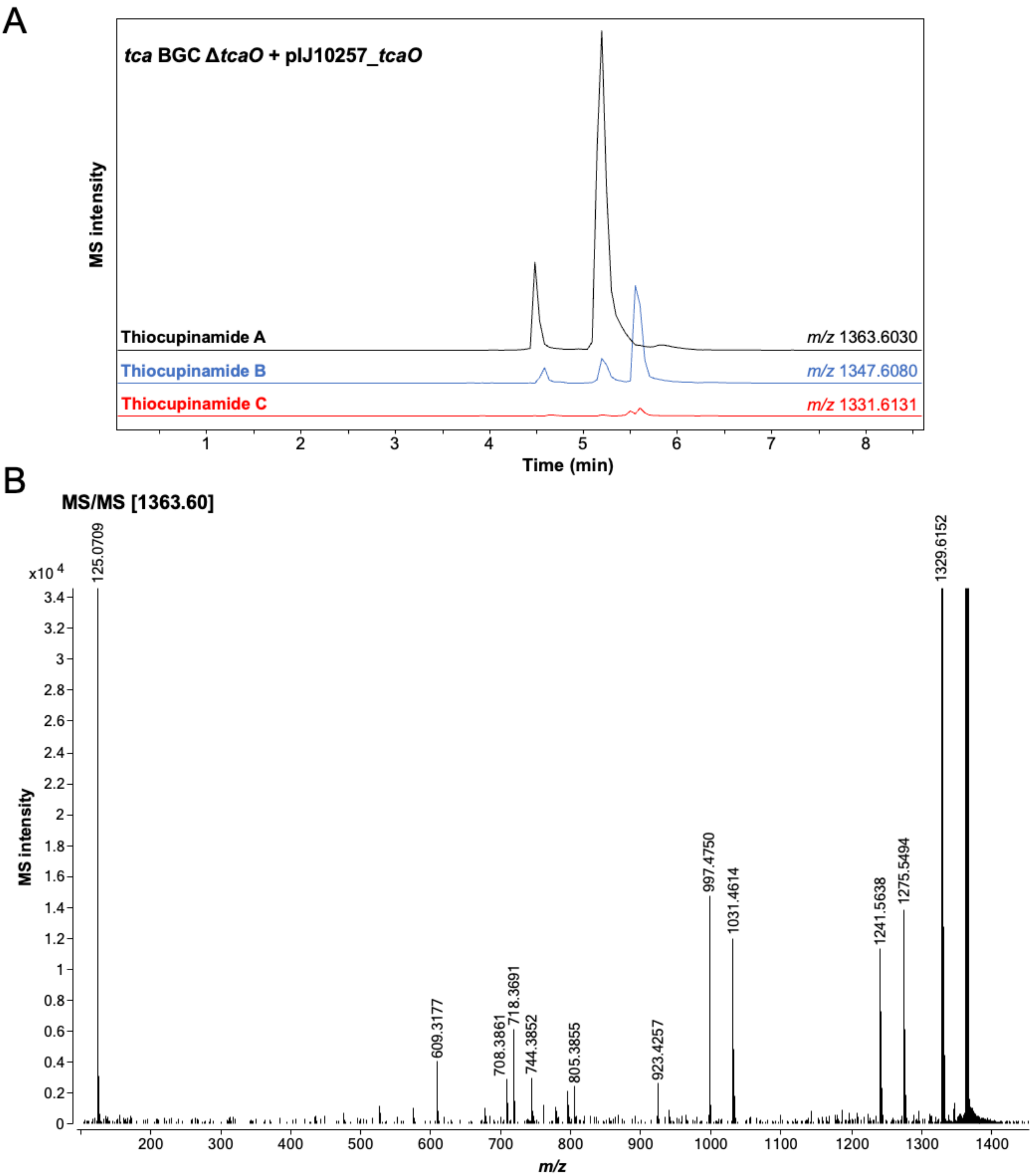


**Figure S7** Complementation of Δ*tcaO* mutant with pIJ10257_*tcaO*. **A.** Extracted ion chromatograms showing the relative proportion of the molecules detected in *S. coelicolor* M1146 integrated with pSET152 P*tsrA tca* BGC Δ*tcaO* and pIJ10257_*tcaO*. Extracted ion chromatograms are shown for each m/z (10 ppm extraction window) and are scaled relative to the largest overall peak. **B.** MS/MS spectrum of thiocupinamide A obtained on an Agilent G6546A Q-TOF LC-MS. The spectrum has been magnified to view fragment details, where each peak matches to fragments characterised in Figure S8.

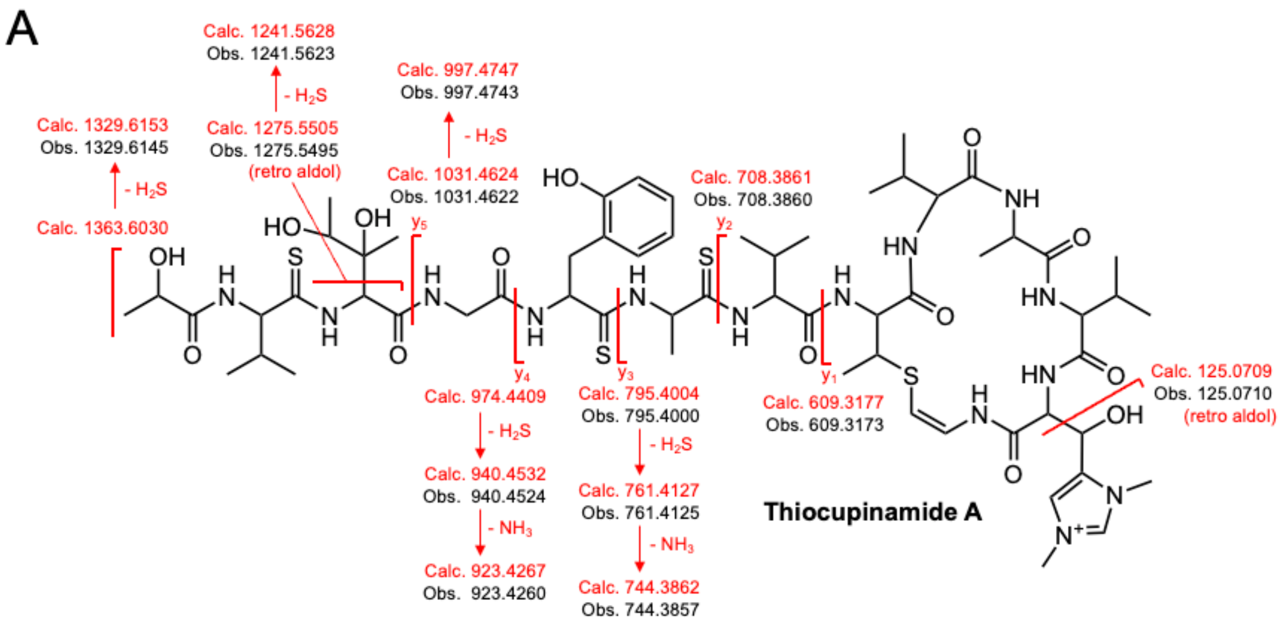


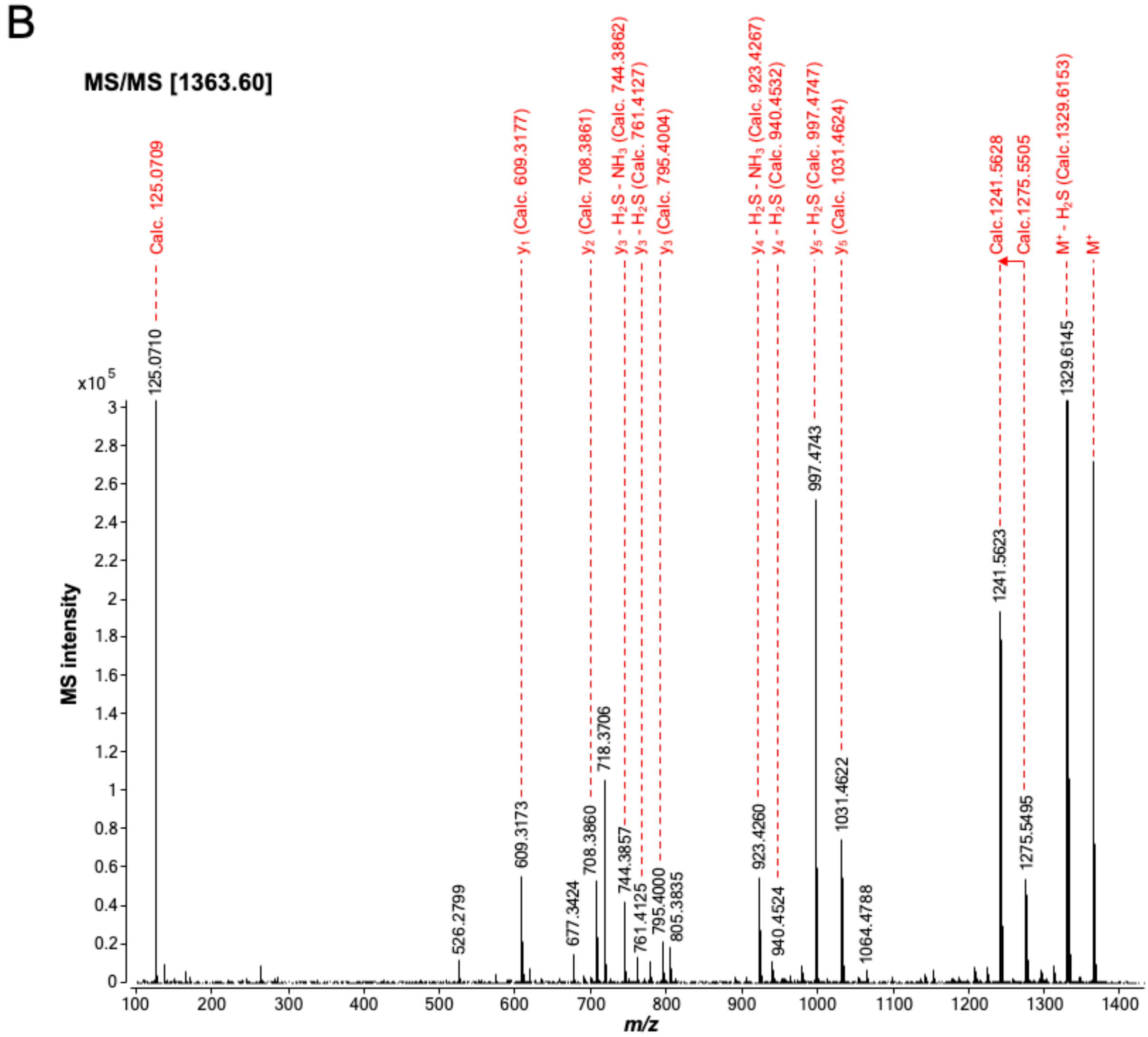


**Figure S8** MS/MS analysis of thiocupinamide A. **A.** Calculated and observed fragments of the predicted structure of thiocupinamide A. **B.** MS/MS spectrum obtained on an Agilent G6546A Q-TOF LC-MS with a collision energy of 70%. The spectrum has been magnified to view fragment details.

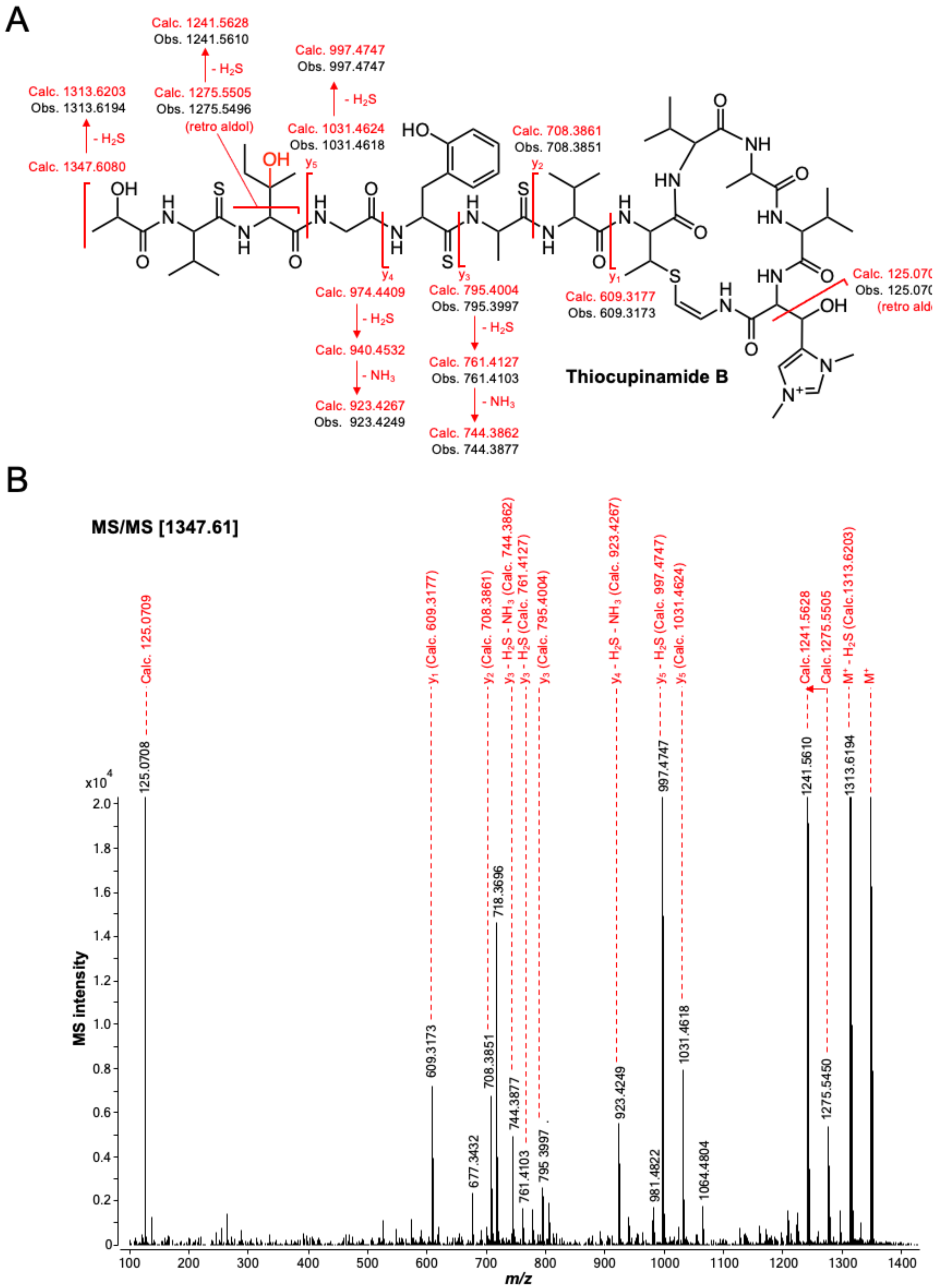


**Figure S9** MS/MS analysis of thiocupinamide B. **A.** Calculated and observed fragments of the predicted structure of thiocupinamide B. The location of the single hydroxylation is predicted based on the presence of fragments consistent with a retro aldol fragmentation of β-hydroxylated Ile. **B.** MS/MS spectrum obtained on an Agilent G6546A Q-TOF LC-MS with a collision energy of 70%. The spectrum has been magnified to view fragment details.

A

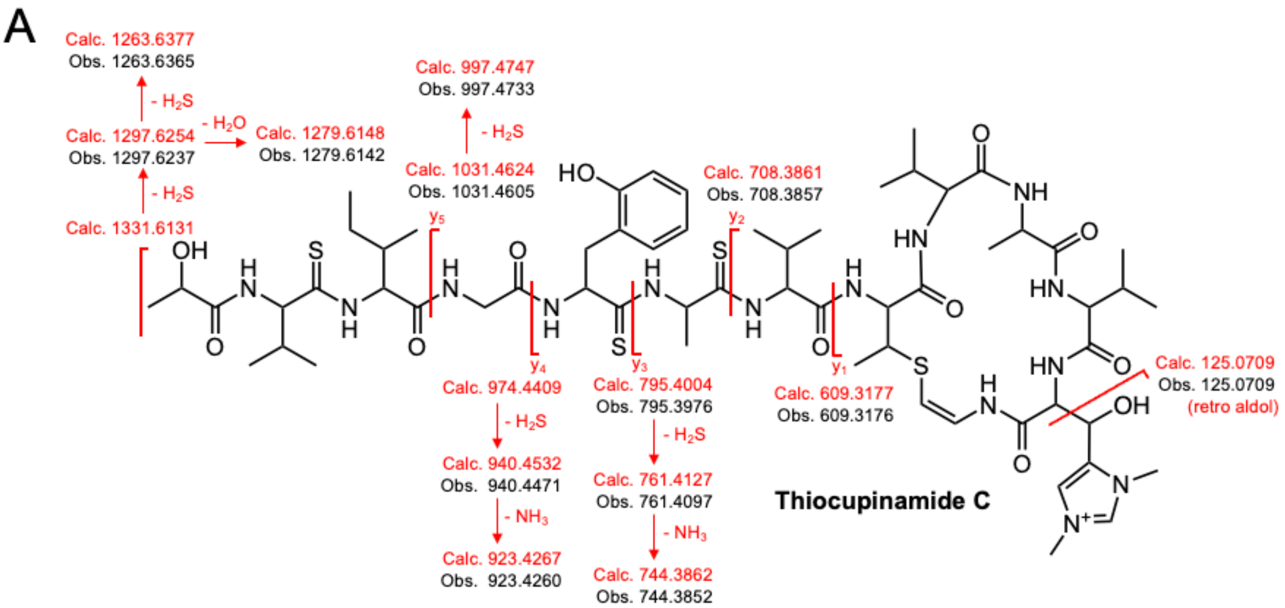


B

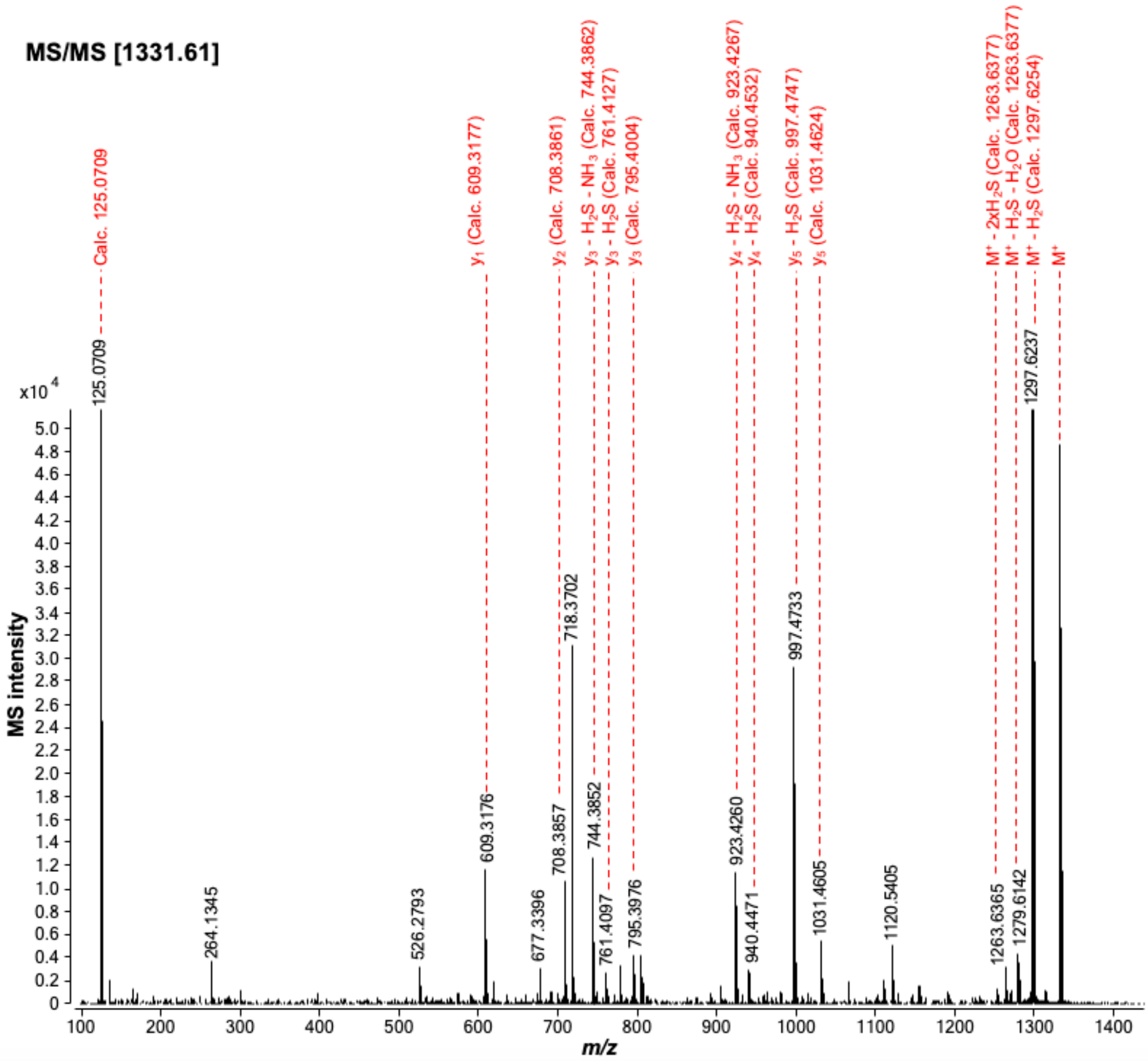


**Figure S10** MS/MS analysis of thiocupinamide C. **A.** Calculated and observed fragments of the predicted structure of thiocupinamide C. **B.** MS/MS spectrum obtained on an Agilent G6546A Q-TOF LC-MS with a collision energy of 70%. The spectrum has been magnified to view fragment details.

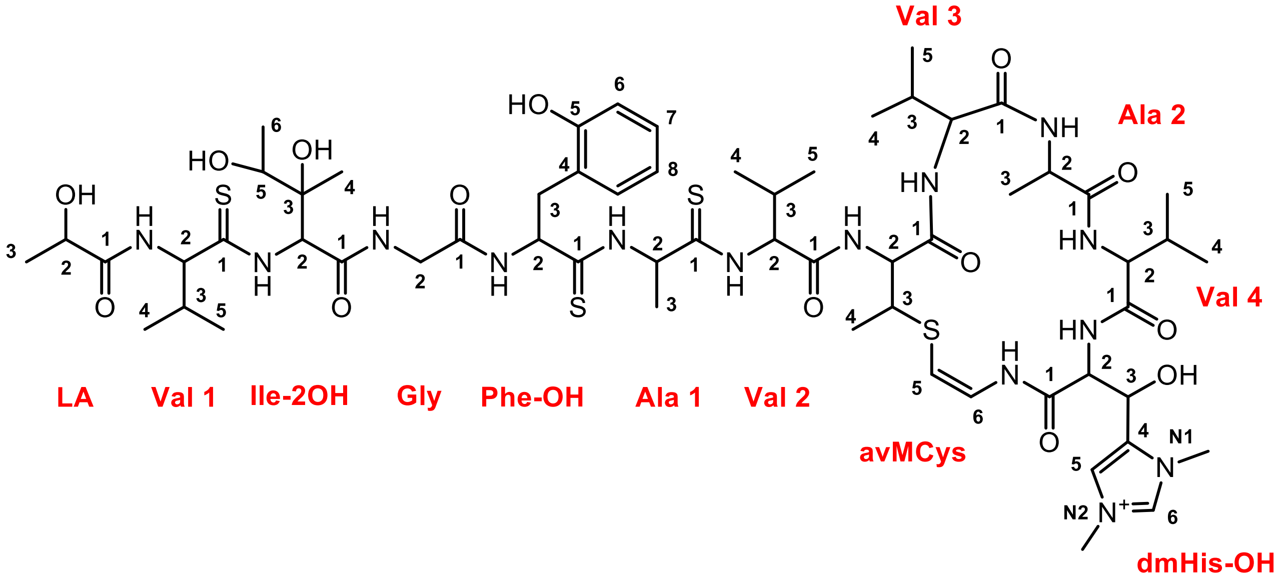


**Figure S11** Thiocupinamide A residue naming and numbering scheme used in Table S5.

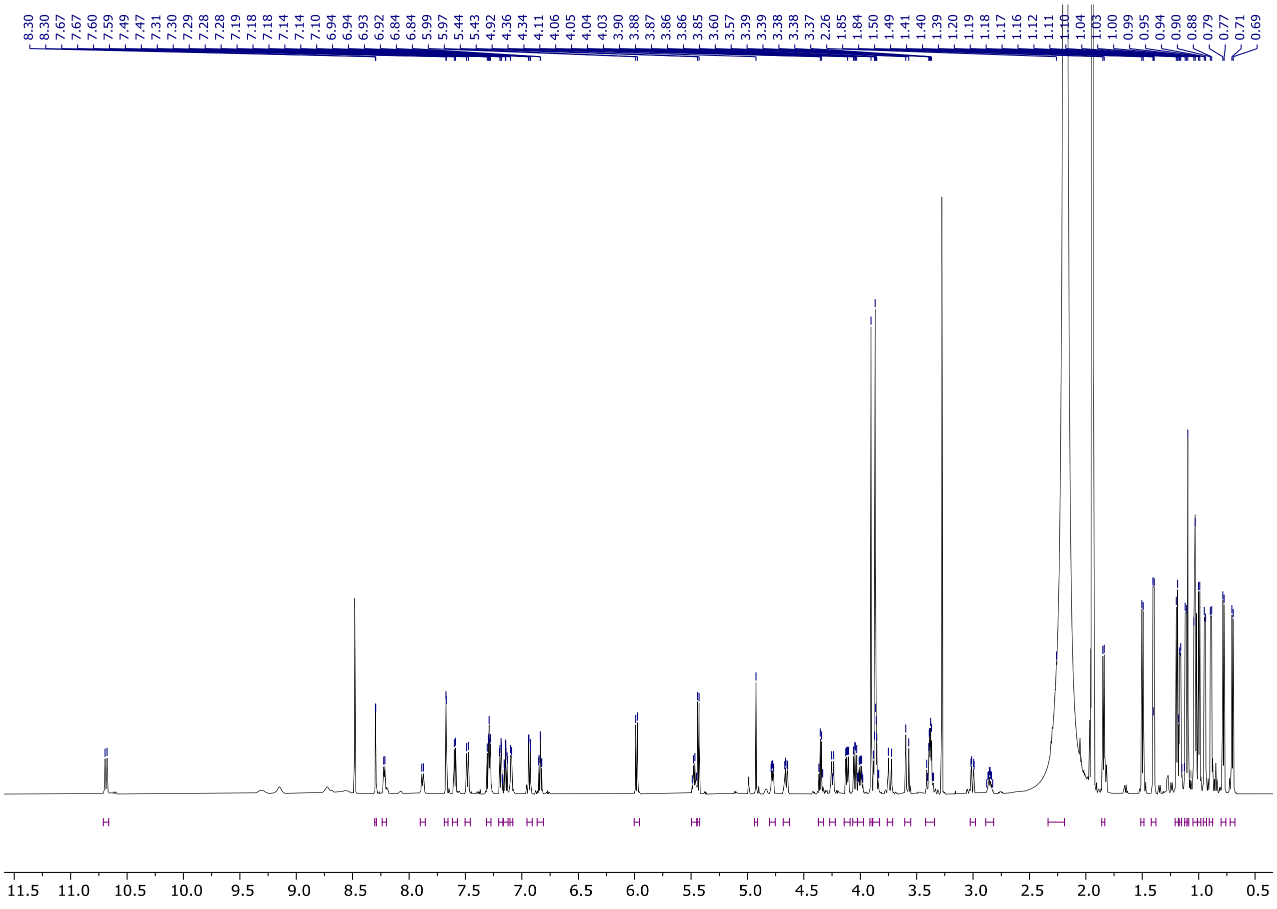

**Figure S12** Thiocupinamide A $^{1}H$ NMR spectrum (600 MHz, acetonitrile-d3, 298 K).

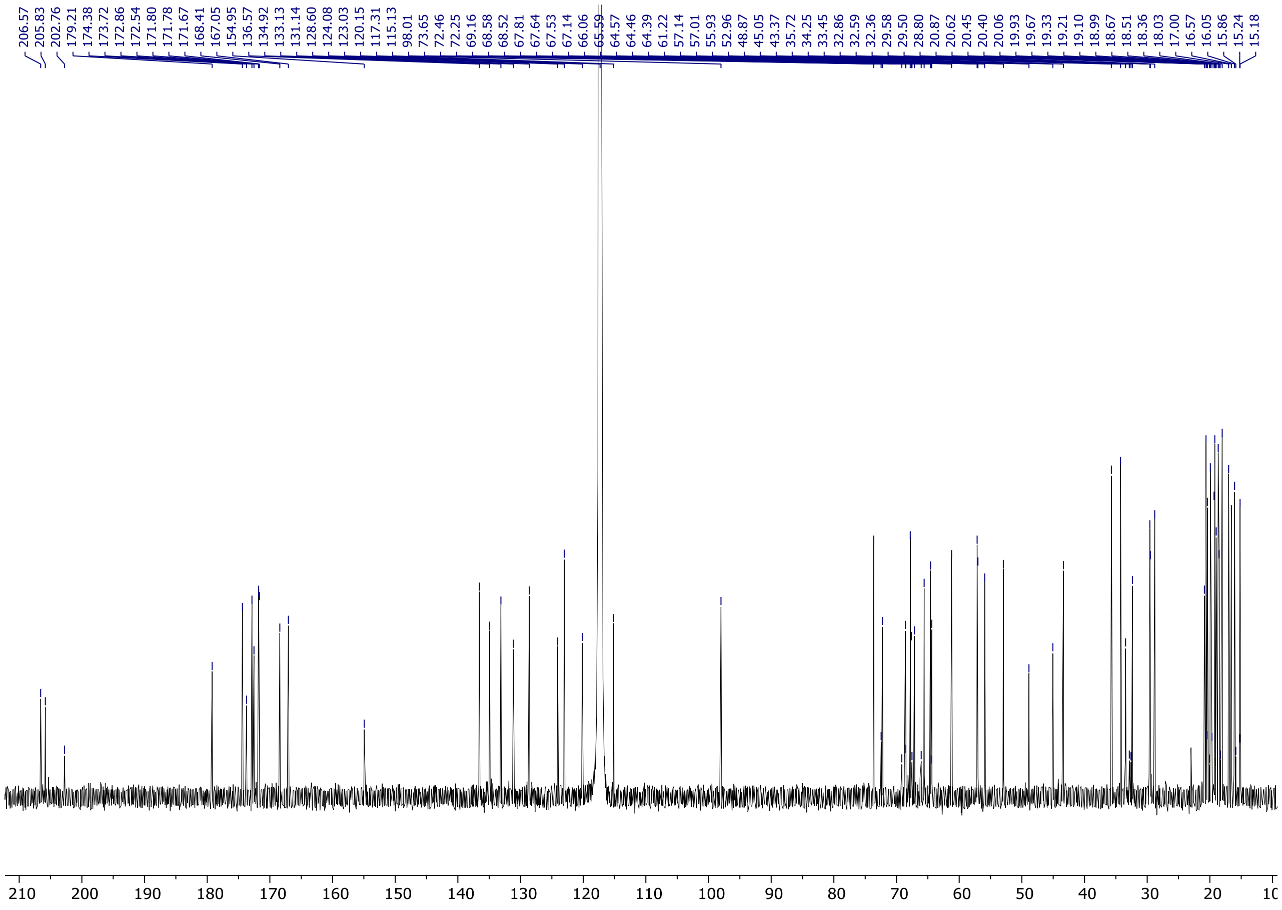


**Figure S13** Thiocupinamide A $^{13}C$ NMR spectrum (150 MHz, acetonitrile-d3, 298 K).

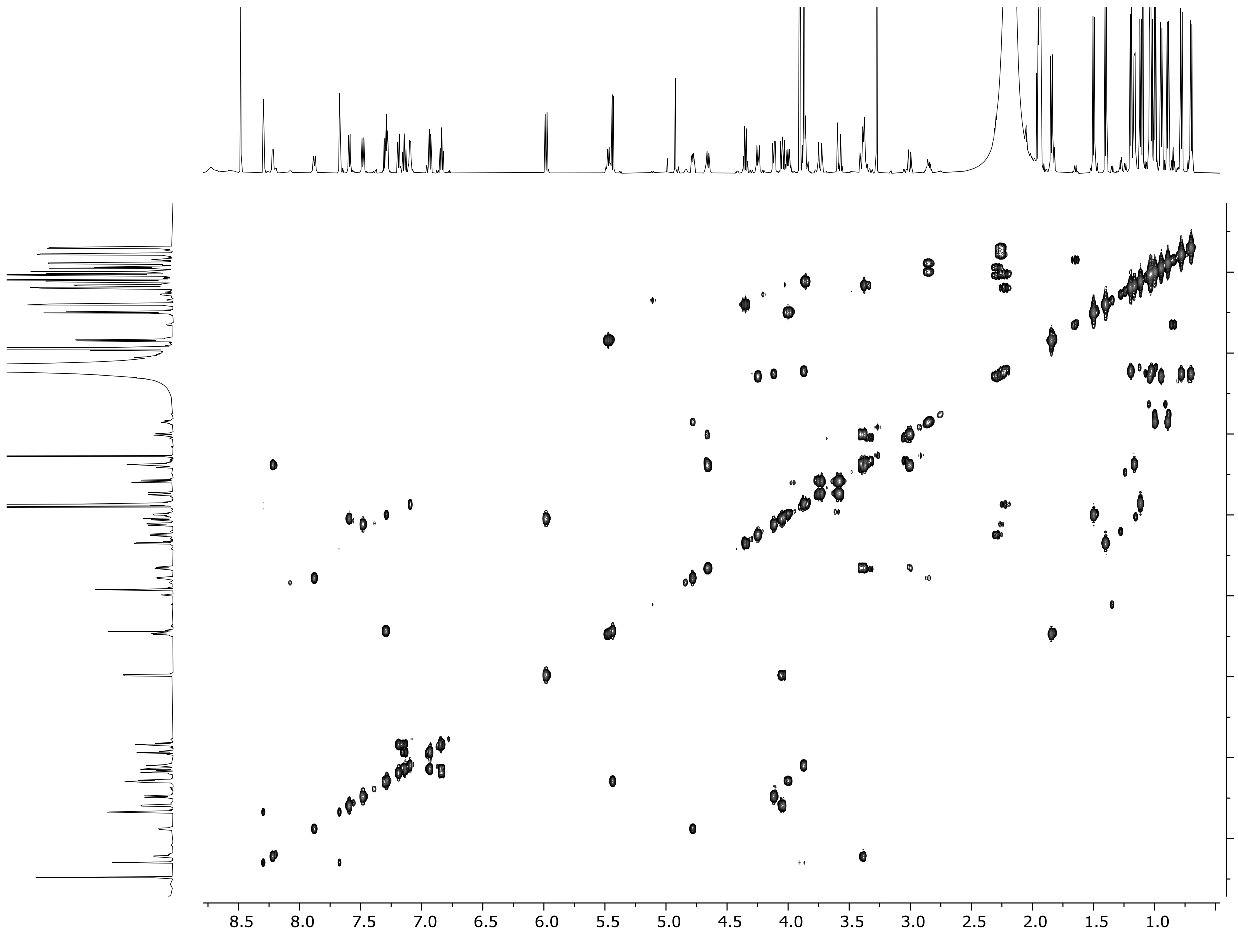


**Figure S14** Thiocupinamide A COSY NMR spectrum (acetonitrile-d3, 298 K).

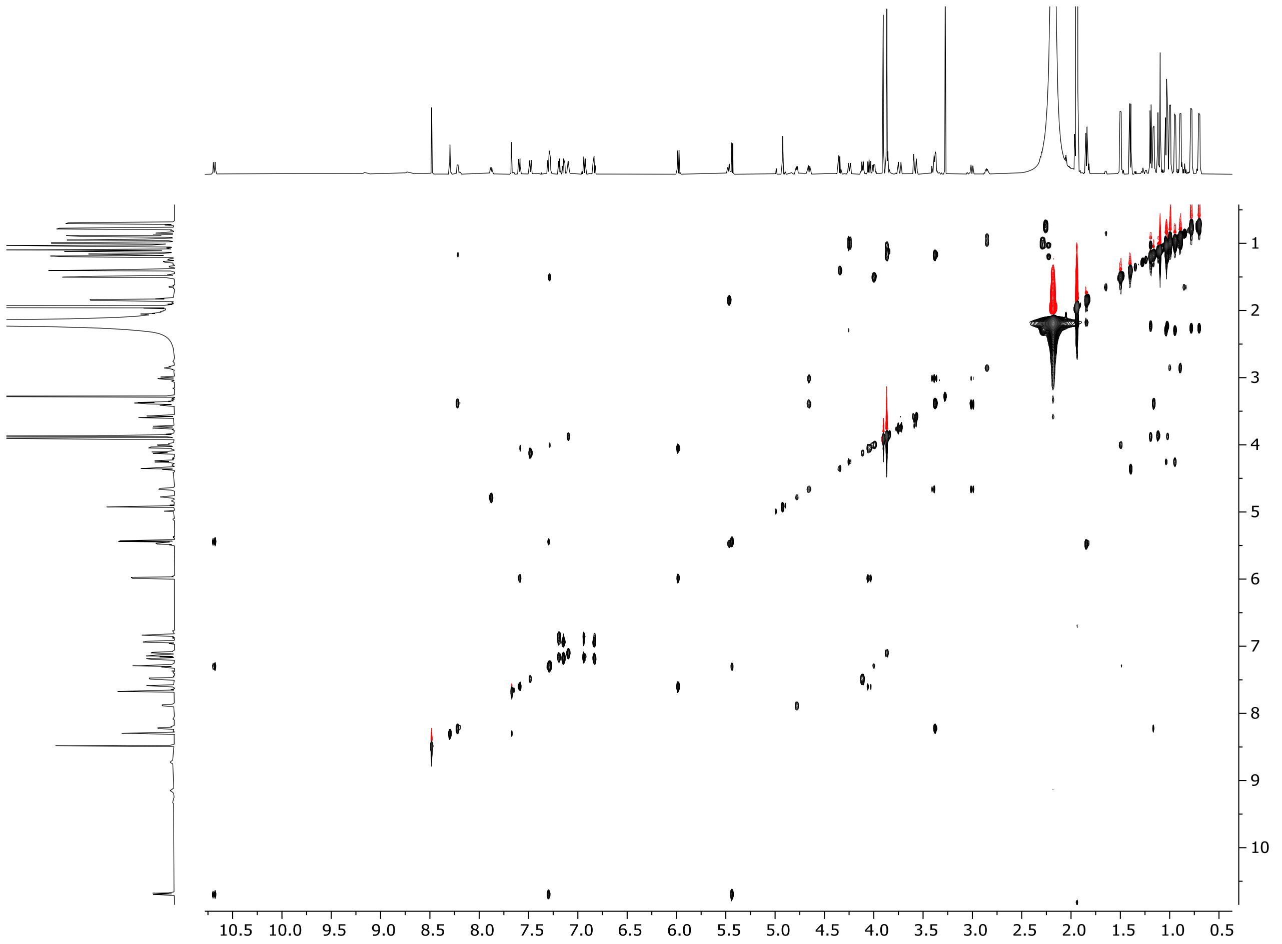


**Figure S15** Thiocupinamide A TOCSY NMR spectrum (acetonitrile-d3, 298 K).

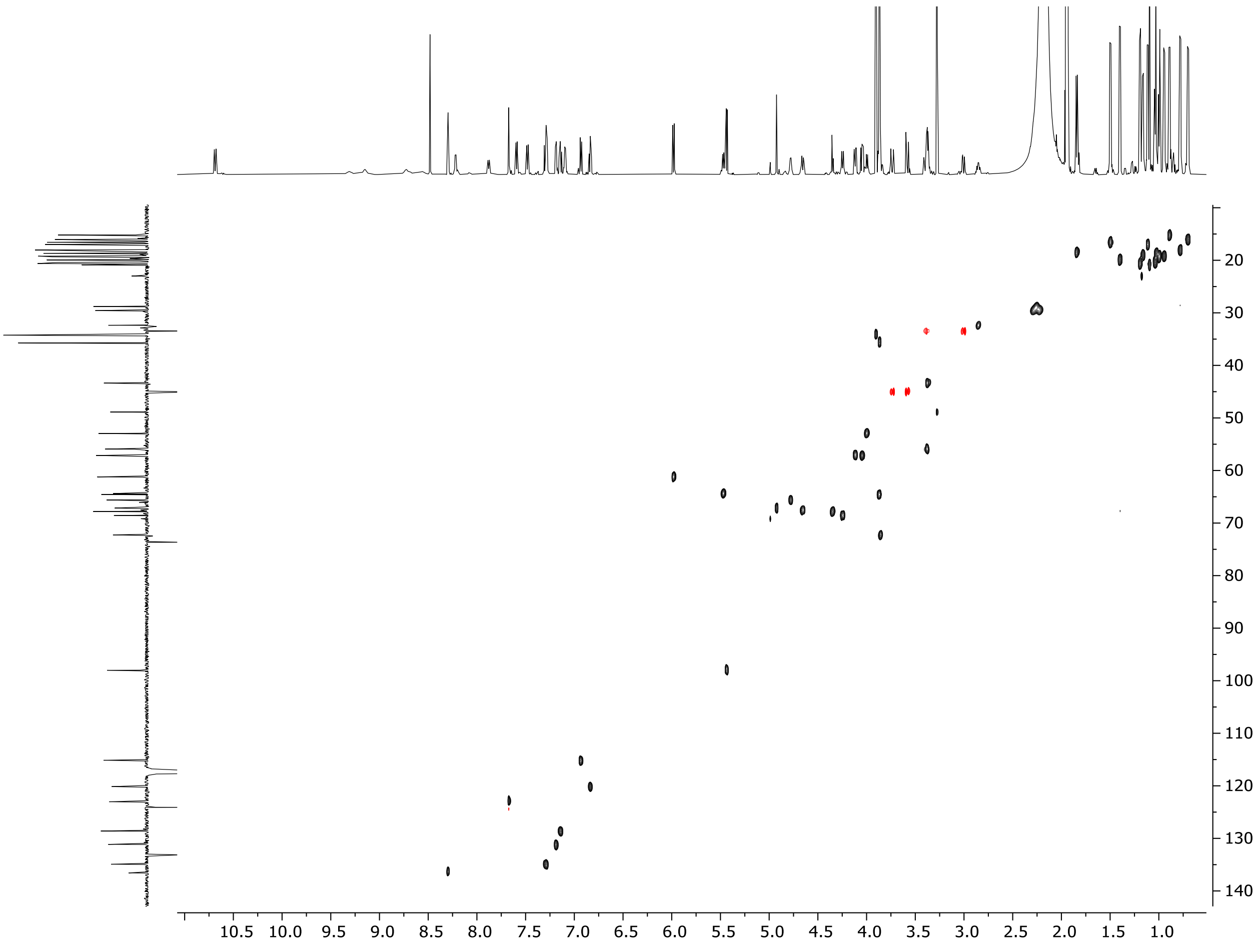


**Figure S16** Thiocupinamide A multiplicity-edited HSQC NMR spectrum (acetonitrile-d3, 298 K).

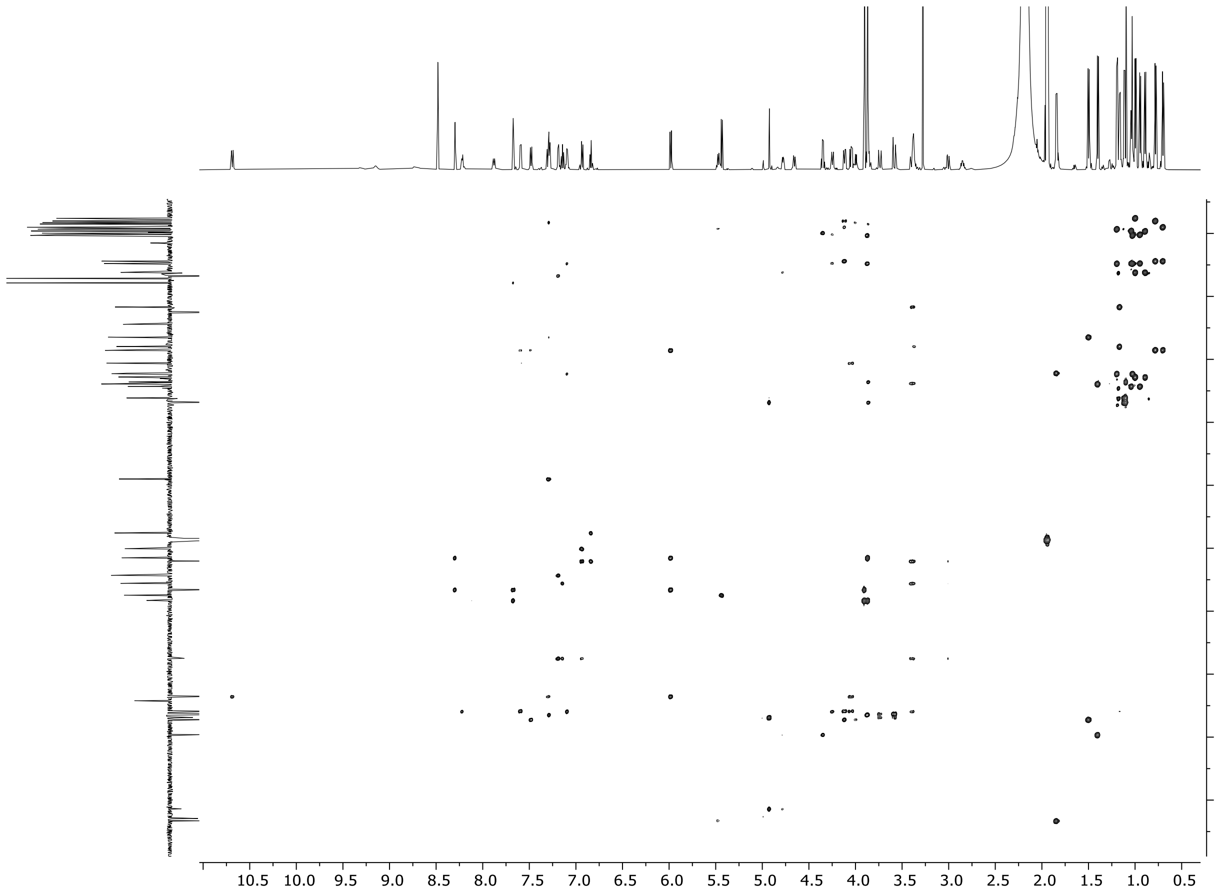


**Figure S17** Thiocupinamide A $^{1}H$-$^{13}C$ HMBC NMR spectrum (acetonitrile-d3, 298 K).

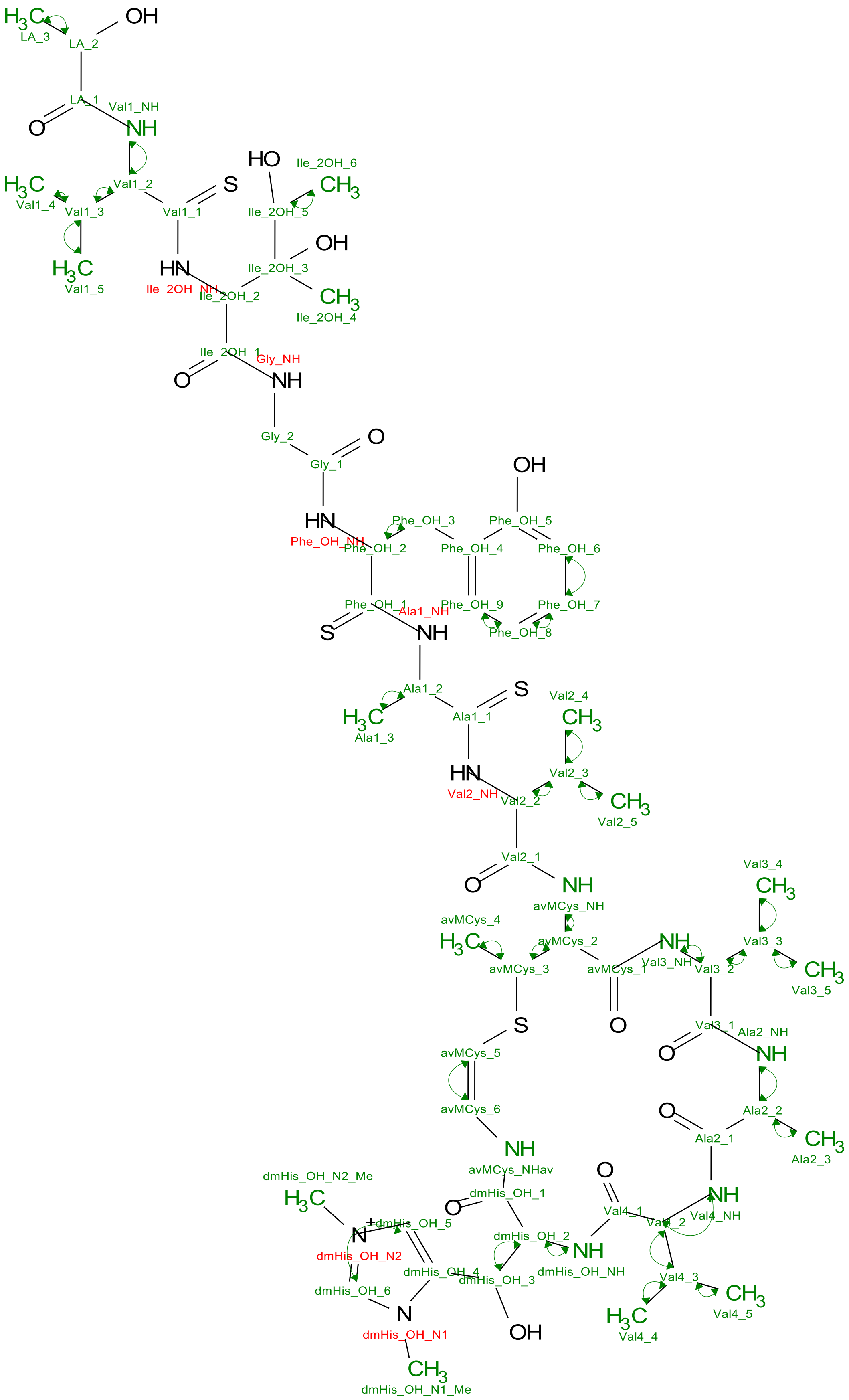

**Figure S18** COSY correlations observed for thiocupinamide A.

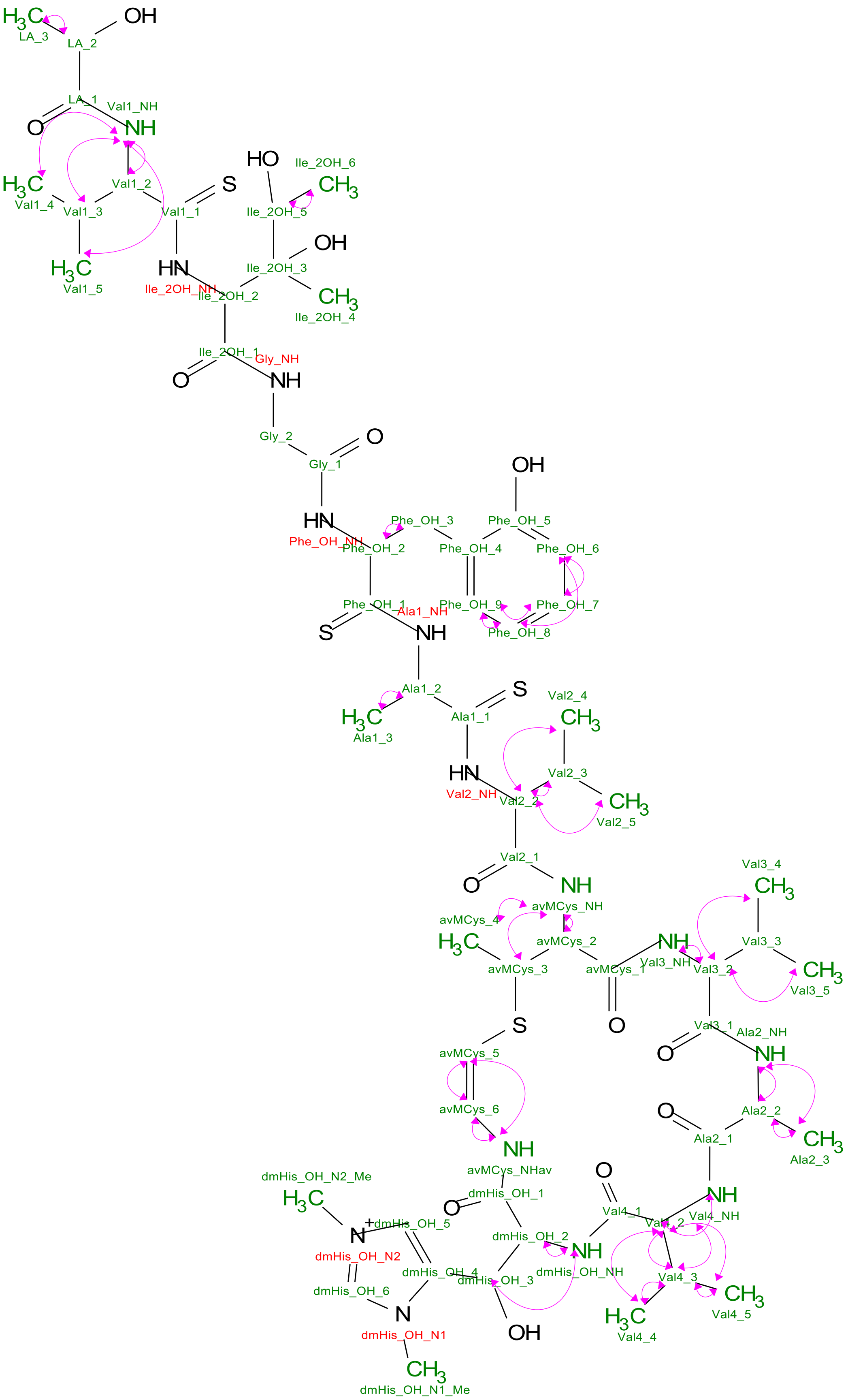


**Figure S19** TOCSY correlations observed for thiocupinamide A.

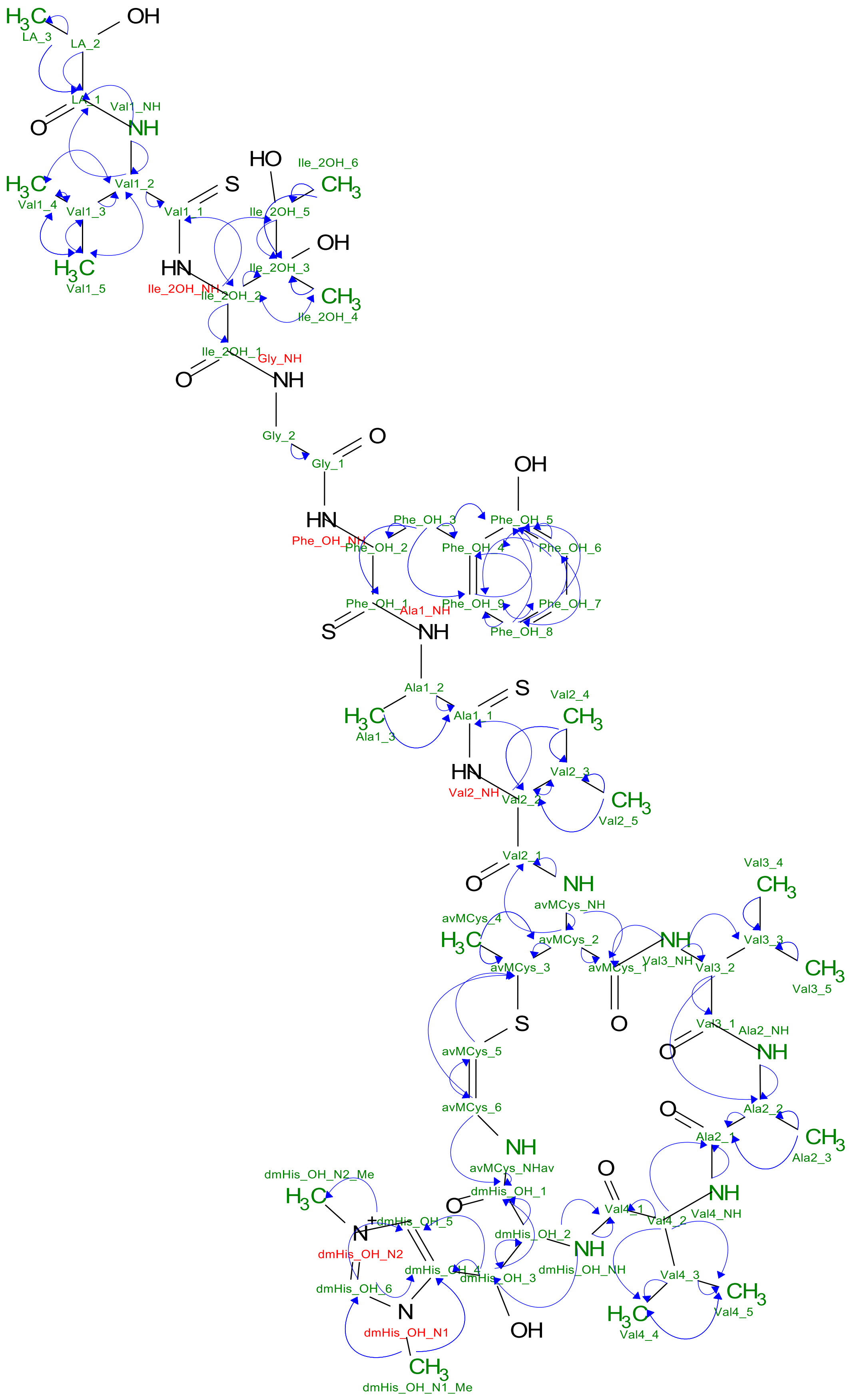


**Figure S20** $^{1}$H-$^{13}$C HMBC correlations observed for thiocupinamide A.

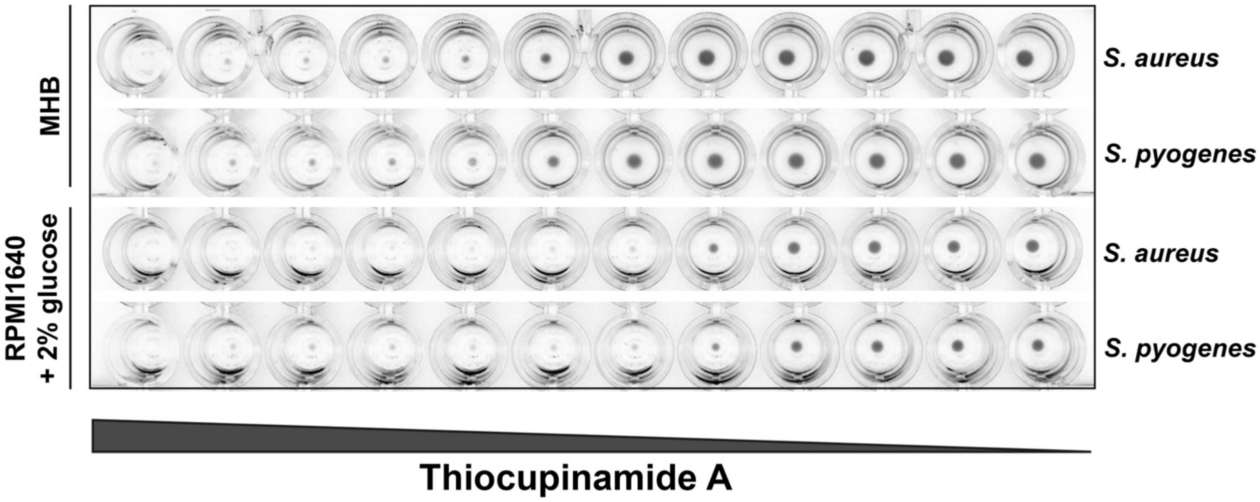

**Figure S21** Thiocupinamide A inhibitory activity on *S. aureus* and *S. pyogenes* growth, in Muller Hinton broth (MHB) and in RPMI1640 supplemented with 2% glucose. Serial dilution of the compound ranging from 64 to 0.0625 µg/mL were tested.
